# Optimising expression quantitative trait locus mapping workflows for single-cell studies

**DOI:** 10.1101/2021.01.20.427401

**Authors:** Anna S.E. Cuomo, Giordano Alvari, Christina B. Azodi, single-cell eQTLGen consortium, Davis J. McCarthy, Marc Jan Bonder

## Abstract

Single-cell RNA-sequencing (scRNA-seq) has enabled the unbiased, high-throughput quantification of gene expression specific to cell types and states. With the cost of scRNA-seq decreasing and techniques for sample multiplexing improving, population-scale scRNA-seq, and thus single-cell expression quantitative trait locus (sc-eQTL) mapping, is increasingly feasible. Mapping of sc-eQTL provides additional resolution to study the regulatory role of common genetic variants on gene expression across a plethora of cell types and states, and promises to improve our understanding of genetic regulation across tissues in both health and disease. While previously established methods for bulk eQTL mapping can, in principle, be applied to sc-eQTL mapping, there are a number of open questions about how best to process scRNA-seq data and adapt bulk methods to optimise sc-eQTL mapping. Here, we evaluate the role of different normalisation and aggregation strategies, covariate adjustment techniques, and multiple testing correction methods to establish best practice guidelines. We use both real and simulated datasets across single-cell technologies to systematically assess the impact of these different statistical approaches and provide recommendations for future single-cell eQTL studies that can yield up to twice as many eQTL discoveries as default approaches ported from bulk studies.

## Introduction

Expression quantitative trait locus (eQTL, see **Table 1a**) mapping is an established tool for identifying genetic variants that play a regulatory role in gene expression. The approach has been widely applied to bulk RNA sequencing profiles from primary human tissues [1] and cell lines [2], [3], as well as sorted cell populations, e.g. blood cell types [4]. Statistical methods for (bulk) eQTL mapping have been extensively tested over the years, with key findings including the need to control for population structure and covariates [5] and to account for multiple testing to control the false discovery rate [6]. Linear mixed models (LMMs) in particular have become a popular framework for genetic analyses of molecular traits, due to their flexibility and ability to robustly control for confounding factors.

**Table 1.**
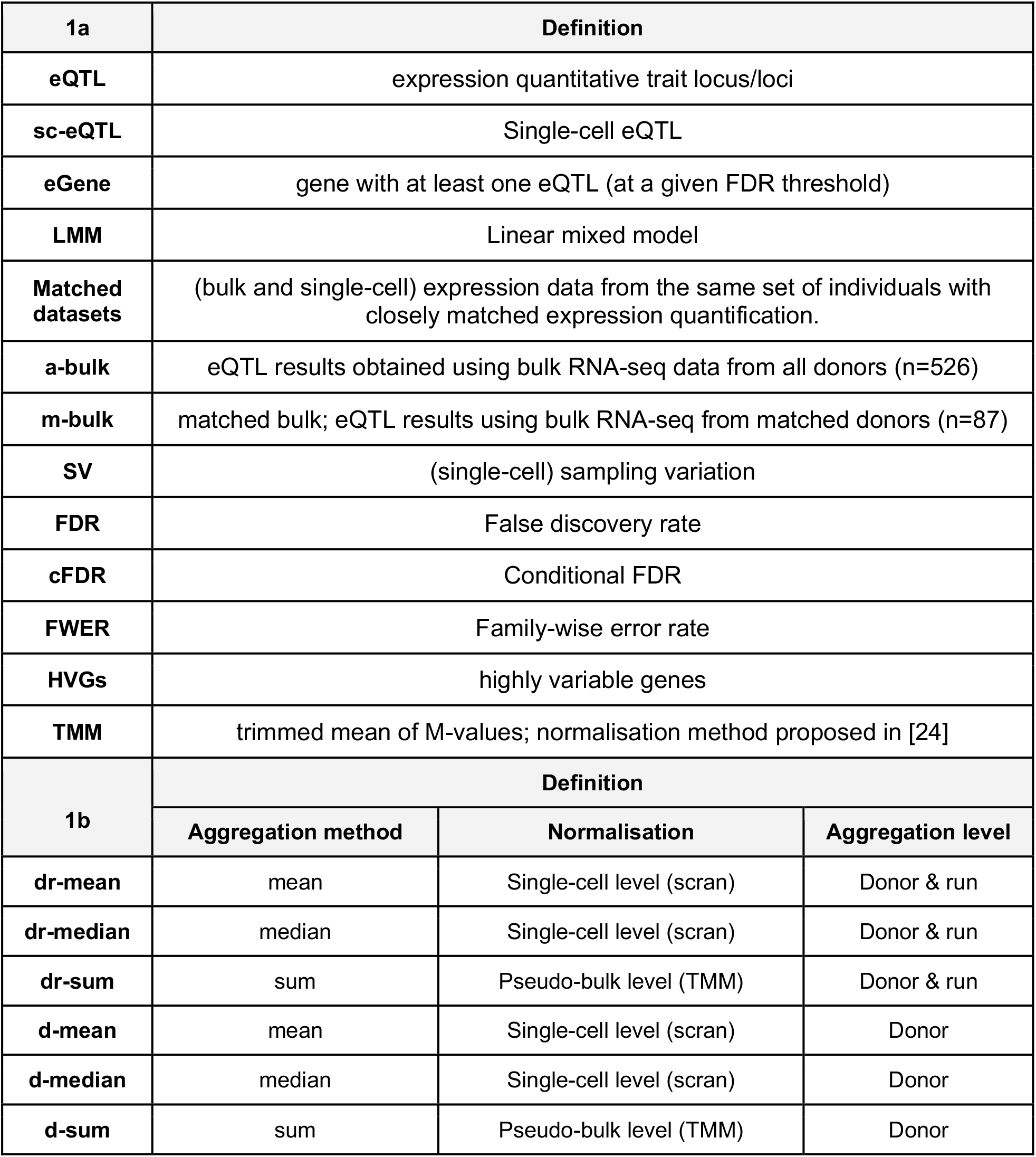
**(a)** Abbreviations used in the manuscript. **(b)** Summary of the six key aggregation-normalisation strategies used in this study. In particular, for each approach we specify the aggregation method used, the type of normalisation adopted, and the level of aggregation selected.

Recent technological advances have allowed molecular phenotypes, including gene expression, to be assayed at the level of single cells. In particular, single-cell RNA sequencing (scRNA-seq) is now an established technique, and can be deployed at population scale, across many individuals, by exploiting multiplexed experimental designs and using appropriate demultiplexing tools [7–9]. The ability to identify cell types and cell states in an unbiased manner from scRNA-seq data from a single experiment can be used to define homogeneous cell populations, quantify expression levels within them, and then map eQTL in each of them separately. As a consequence, studies where single-cell expression profiles (rather than bulk) are used to perform eQTL mapping have emerged recently [9–15], and promise to greatly improve our understanding of the genetic architecture of gene regulation across tissues, in both human disease and development [16].

As scRNA-seq in large sample sizes becomes feasible, it is important to establish ‘best practices’ and benchmark approaches for the design and analysis of genetic studies using single-cell data. Current efforts in this space have primarily focused on the experimental design of single-cell eQTL (sc-eQTL, **Table 1a**) studies, assessing trade-offs between sequencing depth, the number of donors, and the cell count per donor [17, 18]. Such studies conclude that statistical power can be improved on a fixed budget by performing lower-depth sequencing on more cells per sample or on more total samples.

However, the statistical methods needed to adapt bulk methods to map single-cell eQTL have not yet been systematically benchmarked. Notably, several important processing steps need to be performed on single-cell expression profiles, before we can map the effects of genetic variants on them **(Fig. 1a**). First, cell-level gene expression counts can be obtained using a variety of different methods, that are summarised and reviewed elsewhere [19–21]. Second, quality control (QC) steps should be performed at the level of single cells to filter out low-quality cells (see for example [22] for an overview of best practices). Additionally, individual sequencing runs and batches should be examined to determine their overall quality, and poor quality batches should be discarded altogether. Lastly, traditional genetic analysis of gene expression is well-defined within homogeneous cell populations. Thus, cell type assignment needs to be performed prior to genetic mapping. Clustering and cell type assignment algorithms and approaches have been widely described and remain a focus of benchmarking efforts [22, 23].

**Fig. 1.**
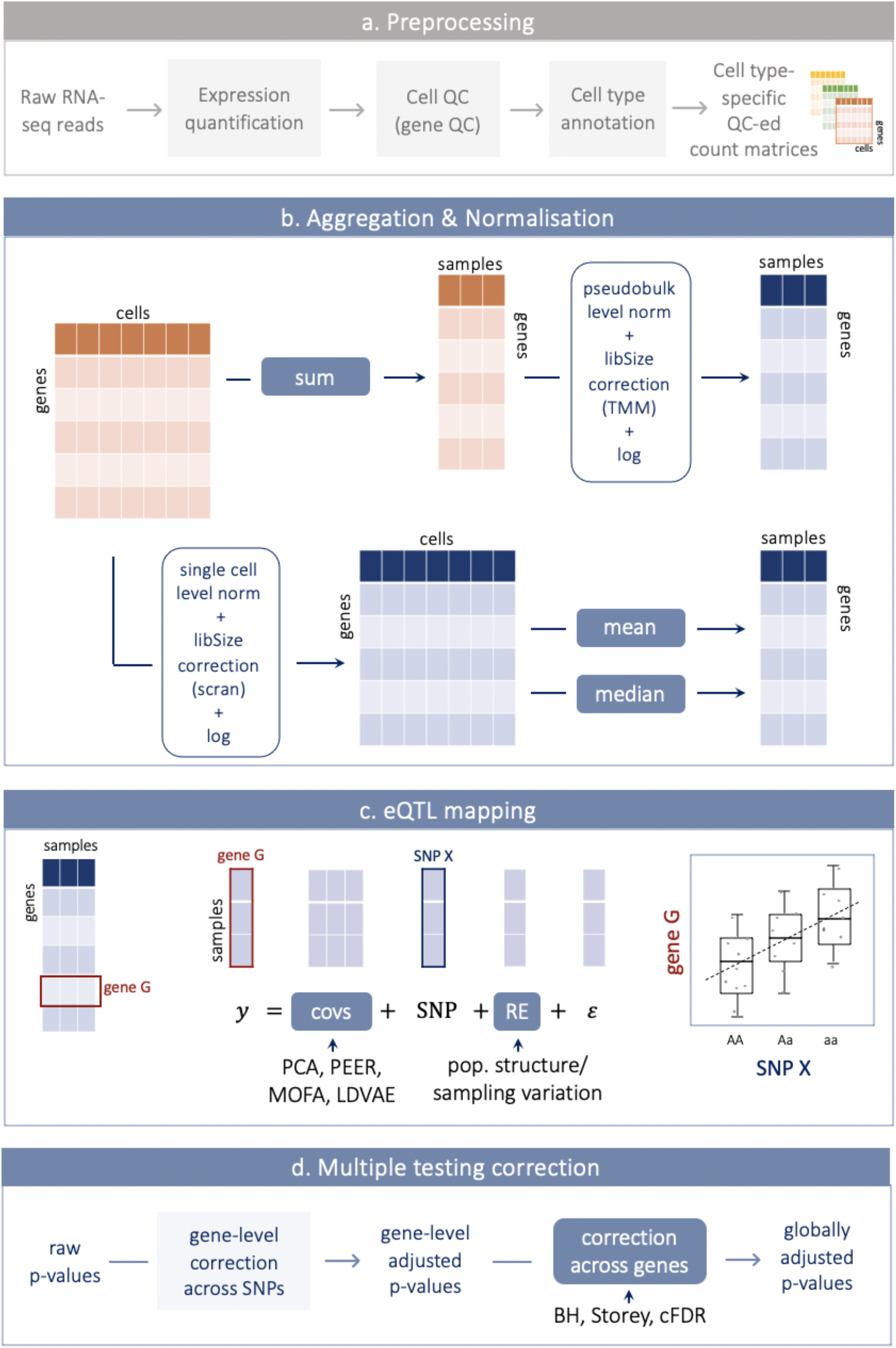
Overview of normalisation, aggregation and the single-cell eQTL mapping considered. **(a)** RNA pre-processing steps to obtain count matrices to perform eQTL mapping, including gene expression quantification, cell and gene-level quality control (QC) and cell type annotation. These steps are not optimised/tested in this work (shown in grey). **(b)** Different approaches tested to perform eQTL mapping using scRNA-seq profiles. Starting from one gene x cell count matrix obtained as in **a**, counts were aggregated per sample (i.e. donor, or donor-run combination), either by summing the data first at sample-level and then normalising using methods designed for bulk RNA-seq (i.e. TMM [24]) as implemented in edgeR) or by first normalising the single-cell counts (using scran/scater [25, 26]) and then calculating the mean or the median at the sample-level. **(c)** eQTL mapping (cis). We map eQTL independently for each gene-SNP pair considered by fitting a linear mixed model. In particular, we model gene expression as the outcome variable (y), the SNP effect as well as additional covariates as fixed effects, and include one (or more) random effect (RE) term to account for population structure and sample variation. We considered various methods to compute covariates, and tested different numbers of covariates as well. (**d**) Multiple testing correction is performed in two steps. First, gene-level p-values are adjusted using a permutation scheme (**Methods**) to control the FWER across SNPs. Second, the top SNP per gene is selected (minimum adjusted p-value; **Methods**) and various methods are used to control the FDR and obtain globally corrected p-values. Steps that we optimise here are highlighted in blue in panels **b, c**, and **d**.

Here, we focused on all aspects downstream of these steps, as these upstream steps are largely dataset- and technology-dependent, and have been previously reviewed and benchmarked. Instead, we set out to optimise statistical analysis workflows that were originally developed for bulk RNA-seq studies in the sc-eQTL mapping setting. In particular, we take advantage of our unique, population-scale, dataset of matched (i.e. from the same individuals) single-cell and bulk expression data from a homogenous cell population (induced pluripotent cells; iPSCs) to evaluate the effect of different statistical choices. Because we have matched samples, we are able to measure performance both in terms of sc-eQTL yield (i.e. the number of significant sc-eQTL) and in terms of concordance with bulk-eQTL, which can be considered as the gold standard for eQTL mapping. We also validate these findings on simulated population-scale scRNA-seq datasets, where the eQTL effects are known, and performance can be measured in terms of power and false discovery rates. Using both empirical and simulated datasets, we identify best practices in terms of aggregation and normalisation, the type and number of expression covariates to include as fixed effects, methods to account for single-cell sampling variation in the model, and methods to guide multiple testing correction using information from bulk RNA sequencing data. Finally, we demonstrate how applying these best practices increases power to detect sc-eQTL and reduce the number of false discoveries.

## Results

### Aggregation and normalisation strategies

Traditional bulk eQTL are germline genetic variants that are associated with differences in gene expression between donors, where the gene expression values represent the summary of a gene’s expression across all cells in the tissue sample. In order to use traditional bulk eQTL mapping methods for single-cell eQTL mapping, we first need to aggregate the multiple measurements (i.e. cells in cluster X or cells of cell-type X) from each donor to obtain bulk-like measurements. Here, we explore different aggregation methods (**Fig 1b**). In particular, we consider the mean, the median, and the sum as aggregation strategies. Initially, we performed aggregation at the donor (“d”) level, i.e. taking all cells for a donor, to maximise the numbers of cells per donor. We call the resulting methods “d-mean”, “d-median”, and “d-sum” (**Table 1b**). Additionally, given that single-cell RNA-sequencing allows for pooling of multiple donors into single sequencing runs (i.e. batches), and for post-hoc grouping of cells (for instance into cell types), we consider aggregating not only at the donor level but also for each individual sequencing run (i.e. all cells from a given donor in a single sequencing run within a single-cell type; designated “dr”, **Table 1b**). While this approach better accounts for variation across technical batches, it also introduces multiple measurements from the same donor. We can account for these repeated measurements in our linear mixed model by including replicate and population structure information as covariates (**Methods**). We call the corresponding methods “dr-mean”, “dr-median”, and “dr-sum”. We chose to explore the dr aggregation level because in the datasets we considered some donors are present in multiple runs [12, 13]. In all cases (i.e. using any of the aggregation methods), aggregated expression values were only calculated for samples (i.e. donors or donor-run combinations) with at least 5 cells.

Importantly, normalisation of the scRNA-seq data was performed in different ways depending on the aggregation method used. For the mean and median aggregation (both at the donor and the donor-run level), we performed single-cell-level normalisation using scran [26] implemented in scater [25], which is one of the standard methods used for single-cell normalisation. The mean and the median were then calculated on the resulting normalised (logged) counts (**Fig. 1b, Table 1**). As an alternative to scran we also tested bayNorm [27] another recent single-cell normalisation approach. We found that the normalised counts are highly correlated (Pearson’s R^2^ mean: 0.88, median 0.93, **Methods**). On the other hand, summed count values (both dr-sum and d-sum) were obtained directly from the raw count data (i.e. non-normalised). Normalisation was then applied on the resulting pseudo-bulk counts, using methods typically used for bulk RNA-seq data. In particular, we perform TMM normalisation on the aggregated counts, as implemented in edgeR [28], one of the best-established methods for bulk RNA-seq normalisation (**Fig. 1b, Table 1**), followed by log transformation.

We test these approaches on two empirical datasets and two simulated datasets. The main analyses are performed on expression data gathered from iPSCs, for which we have bulk RNA-sequencing [29] and Smart-Seq2 single-cell RNA-sequencing [12] data. This unique data allowed us to compare approaches to discovery of eQTL in single-cell expression data, using bulk eQTL results from the same samples as the gold standard for evaluating approaches for single-cell data. Specifically, we assessed the numbers of eGenes discovered in single-cell data and the extent of replication of bulk eQTL effects in single-cell analyses. Additionally, we consider one cell type (midbrain floor plate progenitor; FPP) from a large 10X single-cell RNA-sequencing differentiation study, differentiating iPSCs towards dopaminergic neurons [13]. Lastly, to support our findings, we test our approaches on simulated single-cell eQTL datasets generated using SplatPop (**Methods**), based on either the Smart-Seq2 iPSC expression dataset or the 10X differentiating neuron dataset. In the simulations we could assess eQTL discovery power and the fraction of false positives at a given FDR using the simulated eQTL effects.

### Mapping eQTL using single-cell expression profiles

#### Smart-Seq2 datasets

We first focused on the single-cell data from Cuom*o et al*. [12], from which we selected the iPSC data (day0) to compare to bulk data from the HipSci consortium [2, 29]. We selected the iPSC data because the homogeneous expression profiles of these cells make it an ideal cell type for performing this kind of study. From the HipSci resource we selected 87 healthy donors of European descent for which we had quality controlled: 1) genotype data, 2) Smart-Seq2 sc-RNAseq, and 3) bulk RNA-seq (i.e. matched bulk, from hereon “m-bulk”). Additionally we used a superset of 526 samples (including the previous 87) for which we had quality controlled: 1) genotype data, and 2) bulk RNAseq data, for reference (i.e. all bulk, “a-bulk”). We (re)processed the raw RNA-seq data from the single-cell study to match the bulk processing as much as possible (**Methods**).

We aggregated the single-cell information as described above (**Fig. 1b**) and we tested for *cis*-expression quantitative trait loci (eQTL) using a linear mixed model (LMM) as implemented in LIMIX [30], considering SNPs within 100Kb around the gene and with minor allele frequency (MAF) >10% and Hardy-Weinberg equilibrium P<0.001. We included in the model the first 20 expression principal components (PCs; based on the relevant aggregation method), and used an identical-by-descent kinship matrix to reflect population structure and replicated donors. Each gene’s expression was quantile-normalised prior to being included in the model as phenotype to better suit the assumptions underlying the LMM. We considered the set of 20,545 highly variable genes (HVGs) based on the single-cell measurements (**Methods**). In some instances, to facilitate comparison between the different aggregation/normalisation strategies we selected the set of common HVGs that were tested in every *cis*-eQTL map (n=12,720 genes).

We identified between 776 and 1,835 genes with at least one eQTL (from hereon: “eGenes”, FDR<5%; **Methods**) using the different aggregation methods (out of 12,720 genes tested). To put these numbers in context, the equivalent eQTL map using matched samples with bulk RNA-seq identified 2,590 eGenes (**Table 2**). The large difference in eGene discovery power between the bulk and single-cell methods is at least in part explained by the large difference in the total number of reads per donor and its variability across donors (**Fig. S1**).

**Table 2.** Number of eGenes and replication of eQTL for the different aggregation & normalisation strategies in Smart-Seq2 iPSC cells. The same set of 12,720 genes were considered in all of the strategies. Discovery FDR was controlled at 5% for the discovery; replication was defined as FDR<10% and consistent direction of effect in the two bulk studies, i.e. matched bulk (N=87, m-bulk) and all bulk set (N=526, a-bulk).

|  | Discovery (FDR 5%) |  | m-bulk (FDR 10%) |  | a-bulk (FDR 10%) |  |
| --- | --- | --- | --- | --- | --- | --- |
|  | eGenes | % tested | # replicated | % replicated | # replicated | % replicated |
| **dr-mean** | 1835 | 14.43% | 889 | 48.45% | 1367 | 74.50% |
| **dr-median** | 1337 | 10.51% | 650 | 48.62% | 952 | 71.20% |
| **dr-sum** | 1463 | 11.50% | 819 | 55.98% | 1153 | 78.81% |
| **d-mean** | 1305 | 10.26% | 768 | 58.85% | 1046 | 80.15% |
| **d-median** | 776 | 6.10% | 470 | 60.57% | 625 | 80.54% |
| **d-sum** | 1174 | 9.23% | 709 | 60.39% | 951 | 81.01% |
| **m-bulk** | 2590 | 20.36% | - | - | 2448 | 94.52% |

Overall, we observe two main trends. First, aggregation at the donor-run level outperforms aggregation at the donor level only (e.g. dr-mean vs d-mean). Next, our results indicate that mean aggregation (after single-cell-specific normalisation; 1,835 eGenes) outperforms sum aggregation (followed by bulk-like normalisation; 1,463 eGenes), and median aggregation performs worst in all cases (1,337 eGenes). As well as finding the lowest number of eGenes, the median methods are also responsible for the low intersection of HVGs from 20,545 to 12,720, due to cells with low read counts. When dropping median from the comparison the increase from d-mean to dr-mean is even higher (40% increase in eGenes considering all shared genes vs 46% increase in eGenes considering all tested genes, **Table S1**)). For consistency, we assessed sc-eQTL mapping on the bayNorm [27] dr-mean normalised data and found that the results were very similar compared to the other dr-mean results (1,835 vs 1,702 eGenes, and Pearson’s correlation between p-values R=0.95 and effect sizes R=0.99, p-value < 2.2×10^−16^ in both cases).

Next, we used two selected sets of bulk iPSC RNA-seq data as described above, i.e. m-bulk (n=87) and a-bulk (n=526), to assess the replication of the iPSC sc-eQTL mapping results in bulk data (assumed to be the gold standard, **Table 2, Fig. 2**). We assess replication of the top eQTL effects in a single-cell method in bulk (i.e. direct eQTL replication), and define replication as FDR<10% (in the replication set) and a consistent effect direction. Replication rates from the two sets of samples show a very similar picture: on average we find slightly lower replication rates for the single-cell normalisation methods, but a substantially higher total number of replicated discoveries at the eQTL level. In particular, the highest number of replicated eQTL is found for dr-mean (1,450 considering a-bulk) and highest fraction of replication is found for d-sum (82%, a-bulk). Replication fractions remain consistent when considering all HVGs for eQTL (**Table S1**). Moreover, we see higher replication rates considering a-bulk as compared to m-bulk, indicating that some of the effects found in the single-cell data can only be picked up from bulk datasets with more samples. When specifically looking at the effects that get replicated, we observe that they are highly overlapping (86%) across aggregation strategies. This result indicates that the same effects that get replicated in d-sum, are also replicated in dr-mean, but since there are more effects found in dr-mean the fraction is lower.

**Fig. 2.**
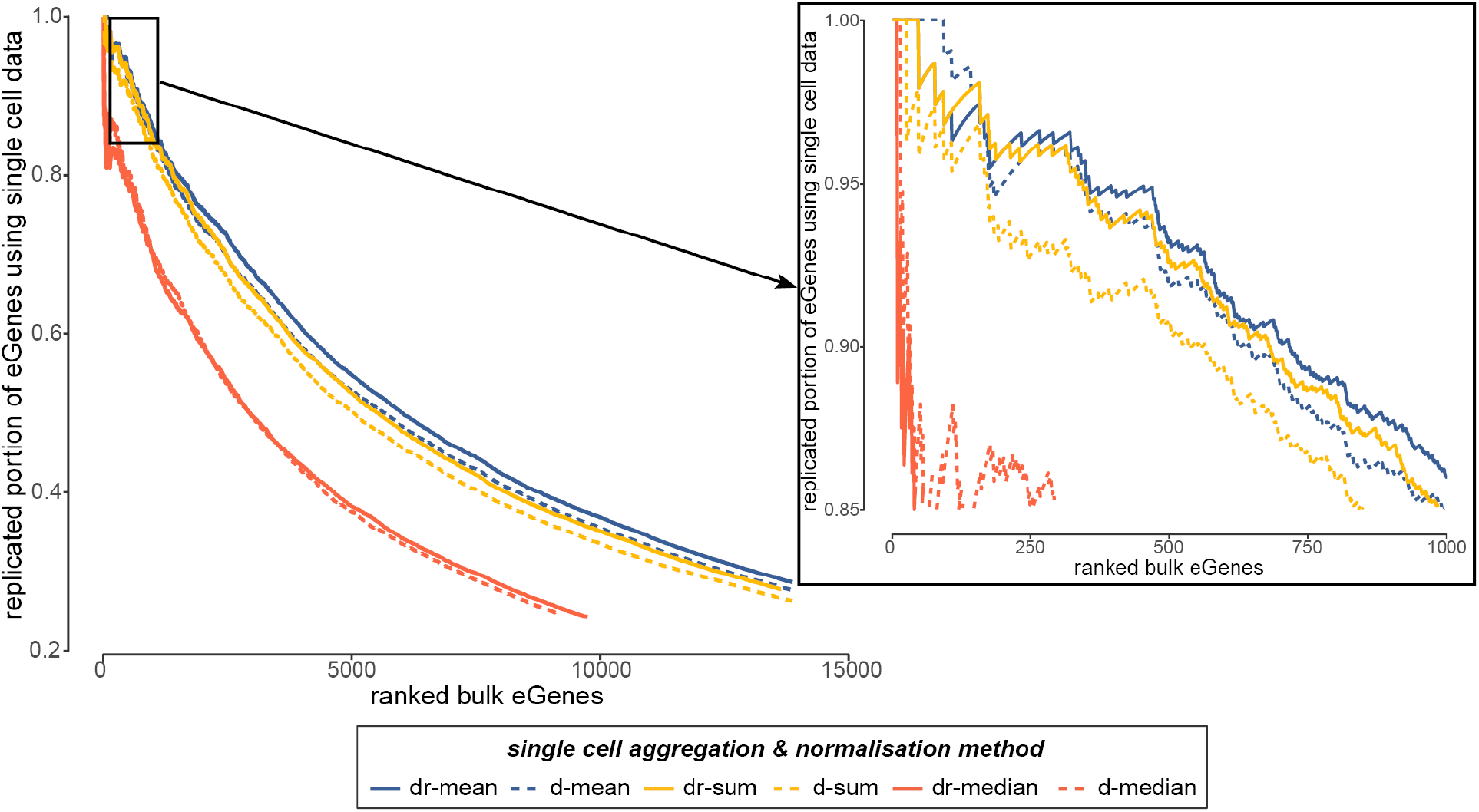
Replication of bulk eQTLs in single cells. Replication rates of bulk eQTL in sc-eQTLs as ranked by eGene significance (p-value) in bulk. Shown are the replication rates of a-bulk eQTL for the six different aggregation/normalisation approaches (mean, median, sum in blue, red, yellow respectively; dr-aggregation level and d-aggregation level in solid and dashed lines respectively).

While the percentage of bulk-eQTL replicated by sc-eQTL mapping in a matched dataset is a powerful performance metric, a limitation of empirical benchmarks like this is that the ground truth is not known. To validate our results on a dataset with completely known true eQTL effects, we simulated single-cell expression profiles for genes on chromosome 2 for a population with known eQTL effects applied to 35% of genes (total expressed genes on chr2: n=1,255, eQTL Genes=439). To ensure the simulations reflected real data, expression statistics from the iPSC Smart-Seq2 dataset were used to estimate key parameters used in the simulations (**Fig. S2**; see **Methods** for details). We then performed aggregation, normalisation, and eQTL mapping as described for the empirical study on 10 replicate simulated datasets and quantified performance in terms of power (fraction of true eQTL detected), empirical FDR (i.e. fraction of false eQTL detected at an FDR < 5%), and effect size correlation (see **Methods** for details). Mean aggregation resulted in greater power of detection than median (paired-t-test; p.adj=1.9×10^−9^) and sum (p.adj=7.3×10^−11^) aggregation, regardless of aggregation level (repeat measures two-way ANOVA; F(2,18)=0.022, p=0.97)(**Fig. 3a**, results **Table S2**; detailed statistical analysis **Table S3**). The aggregation level had a significant effect on the empirical FDR (F(1,9)=15.05, p=0.004)), with donor-run level aggregation resulting in fewer false positive eQTL (paired t-test: p=0.036) (**Fig. 3b**). Finally, we assessed performance in terms of the correlation between the simulated ground truth eQTL effect sizes and the estimated effect sizes (**Fig. 3c**). Here the interaction between aggregation method and level was also not significant (F(2,18)=0.154, p=0.85), but both were significant on their own, with donor-run performing better than donor level aggregation (paired t-test: p=6.43×10^−4^) and mean performing better than median (p.adj=0.041) and sum (p.adj=7.5×10^− 5^). These results corroborate findings from the empirical data analysis showing that mean aggregation after single-cell level normalisation performs better than sum aggregation followed by bulk-normalisation methods, however median aggregation performed better relative to sum aggregation on the simulated data than the empirical data.

**Fig. 3.**
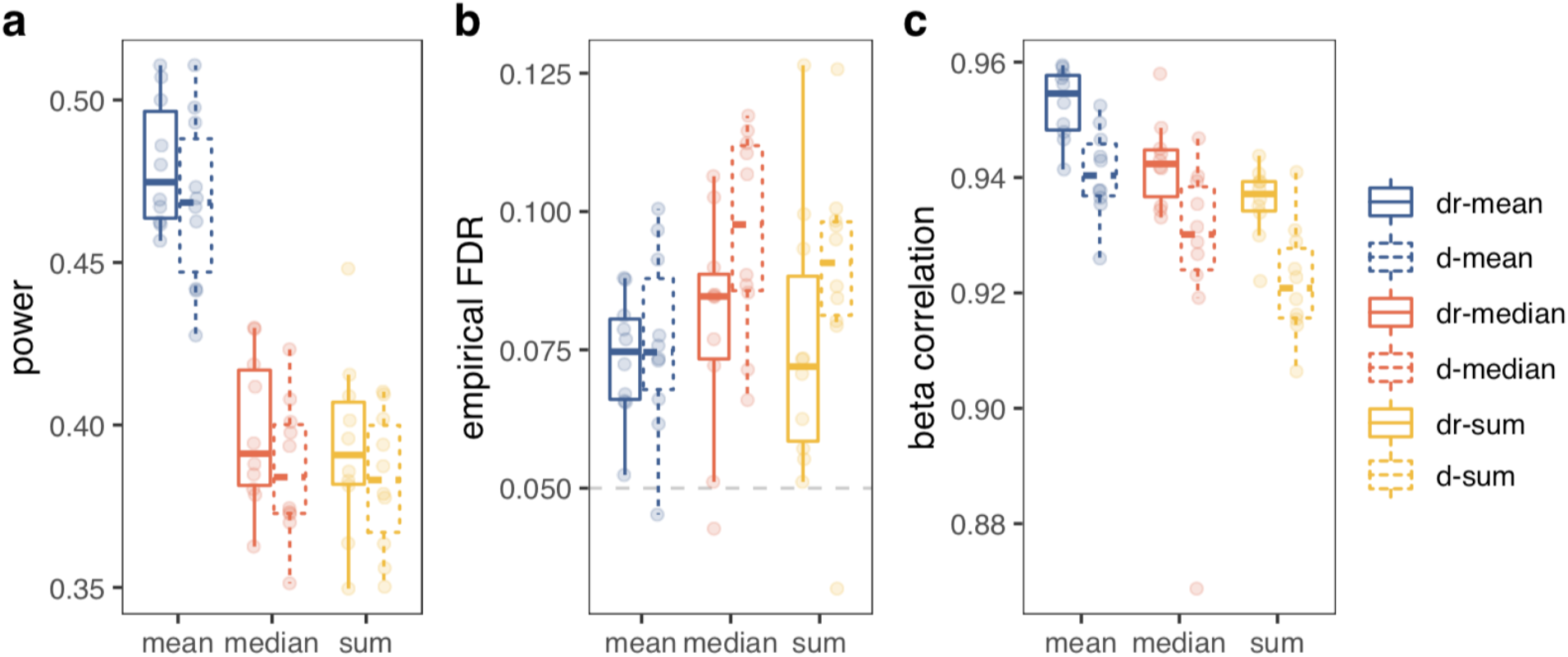
Summary of eQTL mapping performance on simulated Smart-Seq2 iPSC datasets. **(a)** Power to detect simulated eQTL (# true positives / # simulated eQTL). **(b)** Empirical FDR (false discovery rate at FDR < 5%, dashed line). **(c)** Pearson’s correlation between the ground truth and estimated effect sizes for genes simulated as eGenes. Colors and line types are as in Fig. 2. Box plots summarise the distribution, while the points show performance for each replicate (n=10).

A remaining question was if the aggregation level and method would impact eQTL detection power the same way for different sized mapping populations (i.e. number of donors). Using the simulation framework to simulate and map eQTL for populations ranging from 50 to 300 donors, we found that as the number of donors increased the differences in performance between the mean and sum aggregation decreased (**Fig. S3**).

As an alternative to the aggregation-based approaches, we also considered treating individual cells as distinct observations and mapping eQTL directly (after scran normalisation, with no aggregation). We included a random effect term in the LMM to account both for the expected correlation between cells from the same donor and similarity in gene expression between more genomically similar donors (**Methods**). An LMM is often used for “repeated measures” analyses— where multiple measurements are made from the same subject—in preference to classical approaches such as repeated measures ANOVA [31]. We tested this approach on the subset of HVGs on chromosome 2, and considered two distinct approaches: first, we considered all available cells (ranging from 5 to 379 cells per individuals, n=7,552 cells, 2,766 genes tested); second, we subsampled to a fixed number of cells (5 cells for each individual, n=445, 2,718 genes tested) per individual. In both cases, we observed a high number of eGenes (1,668 for the first approach, corresponding to ∼60% genes tested and 840 using the second approach, i.e. 31% of the genes tested). However, the replication rates were very low, ranging from 10 to 15% when considering the two approaches and the two sets of bulk results (**Methods**), likely suggesting inflation and a high rate of false positives, confirmed by poor correlation of effect sizes between these results and those obtained using bulk RNA-seq (**Fig. S4**). To further confirm this result, we tested the first approach on one of the simulated datasets described above. This analysis resulted in a 75.6% false discovery rate, a power of just 18%, and a correlation between the absolute value of simulated and estimated effect sizes of 0.832 (**Fig. S5**), notably worse performance for all three metrics than with aggregation (see above). Given the high false positive rate and the increased computational burden caused by the larger sample sizes when treating each cell as a separate observation, we focus on the aggregation-based approaches for the remainder of this paper.

#### 10X datasets

To check the generality of the results obtained from Smart-Seq2 data, we proceeded to assess the same aggregation/normalisation strategies on data generated with the 10X Chromium technology [32]. Recent large-scale single-cell studies have predominantly used 10X because of the lower cost and higher cell throughput of the droplet-based scRNA-seq technology, as compared to plate-based technologies such as Smart-Seq2 [33]. Though cell throughput is higher, most often the number of reads quantified is lower for 10X studies, and reads are from the ‘3 or ‘5 end of transcripts (or both), but not from full-length transcripts as in Smart-Seq2. Specifically, we selected a midbrain floor plate progenitor (FPP) cell population from a recent differentiation study of iPSCs toward dopaminergic neurons [13] (n=174, **Methods**). Unfortunately, we did not have matching bulk RNA-seq data to assess replication, and thus could only assess differences in terms of *cis*-eQTL discovery power.

In general, we observed very similar effects as found in the Smart-Seq2 dataset (**Table 3**). Again the donor-run methods (dr-mean, dr-sum and dr-median) outperformed donor methods (d-mean, d-sum and d-median), and as for Smart-Seq2 we observe that the single-cell normalisation method with mean aggregation outperformed the sum-based aggregation followed by bulk-like normalisation, supporting the previous observation (both from the simulated and the empirical Smart-Seq2 iPSC data) that single-cell-specific normalisation and treating different runs as separate entities is beneficial. Moreover, for 10X data, single-cell normalisation with median aggregation performed especially poorly, likely due to the greater sparsity in 10X data compared to Smart-Seq2 data. Again, we observe that when considering all HVGs we see a larger increase of eGene discoveries from d-mean to dr-mean (19% increase compared to the initial 16%; **Table S4**).

**Table 3.** Number of eGenes for the different aggregation and normalisation strategies in 10X midbrain floor plate progenitor cells. In total 3,390 genes were considered in all of the strategies, and gene-level FDR was controlled at 5%.

|  | eGenes | % genes tested |
| --- | --- | --- |
| dr-mean | 1545 | 45.58% |
| dr-median | 895 | 26.40% |
| dr-sum | 1064 | 31.39% |
| d-mean | 1310 | 38.64% |
| d-median | 443 | 13.07% |
| d-sum | 764 | 22.54% |

We again validated these results using simulated data based on expression statistics from the neuron differentiation 10X dataset to estimate key parameters used in the simulations (see **Methods** for details). The simulations confirmed that donor-run level aggregation also resulted in a lower empirical FDR on 10X data (paired t-test: p=0.002). They also confirmed that median aggregation performs very poorly on 10X data, however there was no significant difference between mean and sum aggregation on power (p=0.46, **Fig. S6**; see results **Table S2**, see detailed statistical analysis **Table S5**). Consistent with results from the Smart-Seq2 simulations, eQTL mapping on the 10X datasets with a smaller sample size resulted in a greater difference in performance, however here sum aggregation performed better on the smaller datasets (**Fig. S7**).

#### Correcting for global expression covariates

Another important step when mapping eQTL is the correction for batch effects and other known and hidden sources of unwanted variation in expression data [1, 34, 35], hereafter collectively referred to as covariates. In the previous analyses, we used the first 20 PCs as covariates, which is common practice in eQTL studies [34, 35]. Here, we tested the impact of alternative approaches and different numbers of factors to include as fixed-effect covariates in the LMMs used for sc-eQTL mapping. In particular, we compared multiple different methods to capture global expression covariates: probabilistic estimation of expression residuals (PEER; [36]), principal component analysis (PCA), linearly decoded variational autoencoder (LDVAE or linear scVI; [37]), and multi-omic factor analysis (MOFA; [38]), for which we considered two different flavours: with and without sparsity constraints; **Methods**). For each approach we tested the effect of including 5-25 factors as covariates in the LMM, in steps of five. We again considered the iPSC data as a homogeneous cell type for which we have both bulk RNA-seq and single-cell RNA-seq on the same samples available. For these tests we focus on the dr-mean aggregation method, as it performed best in both the simulations and empirical tests.

As previously described, we observed a big increase in the number of eGenes discovered when considering covariates as compared to not considering covariates: the minimum increase is 75% (**Fig. 4**) [36]. However, when comparing the method-specific optimal number of covariates (e.g. 15 for PCA, 25 for PEER, **Fig. 4, Methods**), we observe that both PCA and PEER perform markedly better as compared to the other methods (**Table S6, Fig. 4a**). Both the sparse and non-sparse modes of MOFA perform worse and LDVAE, the only method included that works directly on the single-cell data, produces the smallest increase in eGene discovery. Furthermore, the replication rates of the effects in a-bulk, fixed at 20 PCs as previously used, are similar between the different methods (**Fig. 4b**). Our results also show that more computationally expensive methods such as LDVAE, PEER or MOFA do not perform measurably better than the historical default in bulk eQTL studies of correcting for unwanted variation using principal components.

**Fig. 4.**
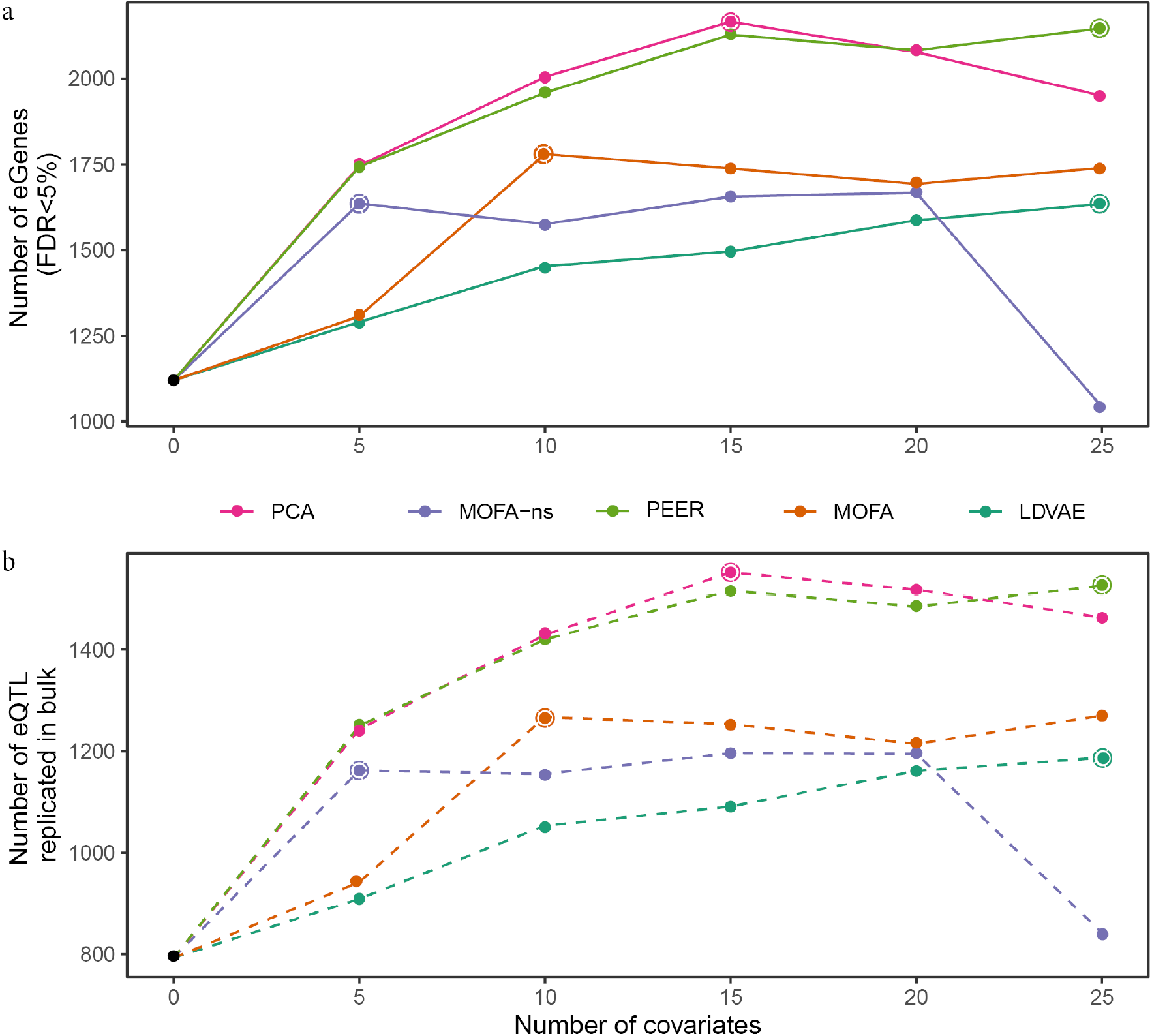
Comparison between covariate-adjustment approaches. **(a)** Number of eGenes obtained when using different approaches to account for covariates, as a function of the number of factors used as covariates (out of 20,545 genes tested). **(b)** Number of replicated discoveries using a-bulk (all samples, n=526). Replication defined as FDR<10% and consistent direction of effect. The optimal number of covariates for each method is highlighted with an open circle.

### Accounting for single-cell sampling variation

After optimising the aggregation-normalisation method and the covariate correction, we moved on to test the effect of accounting for “sampling variation” (SV) in single-cell expression quantifications by including a (second) random effect in our linear mixed model that captures sampling effects (similar to the approach used in [13]; **Methods**). We hypothesised that the total number of reads used to quantify expression per sample and the number of QC-passing cells per sample could contribute to single-cell SV, so we tested the inclusion of random effects defined by these features in the LMM. First, we focused on the d-mean results and replaced the random effect accounting for kinship (baseline) with random effects accounting for either 1/#reads or 1/#cells. Second, we considered dr-mean, which had higher power in eQTL mapping, and tested the use of a joint random effect accounting for both kinship and SV, again either 1/#reads or 1/#cells (in this case we cannot remove the kinship effect term, which reflects the replicated structure of samples across batches, see **Methods** for details).

For d-mean, we observe that replacing the kinship-based random effect with the random effects reflecting SV increases *cis*-eQTL mapping power by >9% (**Table 4**). Additionally, replication rates are similar across all tests, indicating that the additional effects are as likely to be true effects as the initially-discovered eQTL. When considering the dr-mean results, we observe an even stronger increase in eGene discovery power (>21%) when accounting for SV alongside the replicate structure. Again, we observe that the replication rates are comparable to the baseline results (kinship/replicate structure only). For both the d-mean and dr-mean results the 1/#cells random effect seems to work better than 1/#reads.

**Table 4.**
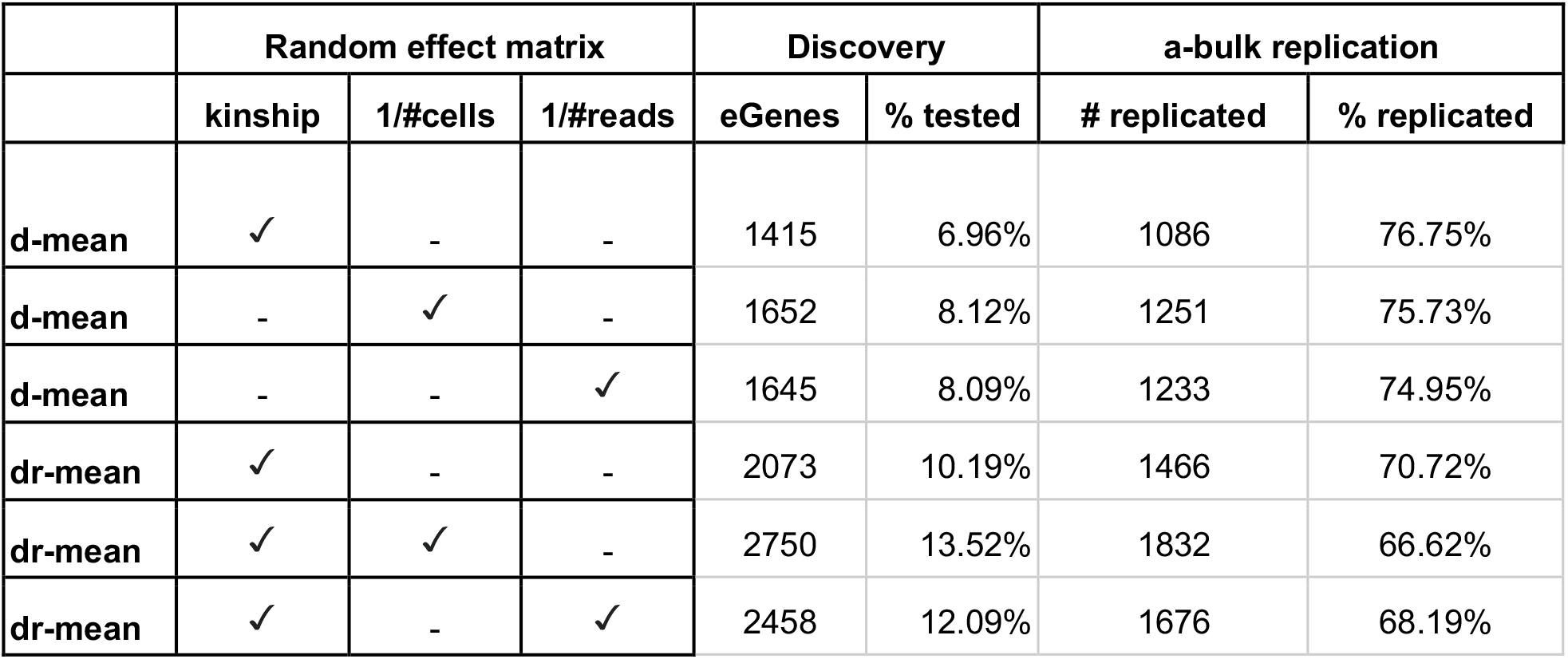
Inclusion of random effect to increase discovery power of sc-eQTL mapping in Smart-Seq2 iPSC sc-RNA seq data. Shown are the number of eGenes that are discovered at an FDR of 5% and the replication in all bulk (a-bulk) defined as FDR <10% and same sign. Tested are the expressed genes on chromosome 2 matched between the two considered aggregations (d-mean and dr-mean, n genes=20,334).

Next, we assessed the impact of the inclusion of the SV measure on eQTL mapping in the 10X dataset. Given the lower read-depths per cell in 10X versus Smart-Seq2, we expected a stronger effect in this setting. However, for the d-mean results we observe a slightly smaller (∼4%) increase in eGene discovery than was observed in the iPSC Smart-Seq2 results (**Table S7**). For the dr-mean based methods, where we include both kinship and SV, we observe again a stronger increase in eGene discovery of around 20%. Like for Smart-Seq2, the SV effect based on 1/#cells allows for the identification of more eGenes than the 1/#reads-based random effect.

### Guided multiple testing increases discovery power

Given that the discovery power of eQTL is heavily dependent on the number of donors for which genetic and expression data is available, and large bulk eQTL studies are available, we set out to leverage bulk data to increase discovery power when mapping sc-*cis*-eQTL. To this end, we selected a recently proposed multiple testing correction method that conditions the false discovery rate (i.e. conditional FDR, or cFDR) on an external set of test statistics [39, 40] and tested the impact of its application to sc-eQTL mapping (**Fig. 5a)**.

**Fig. 5.**
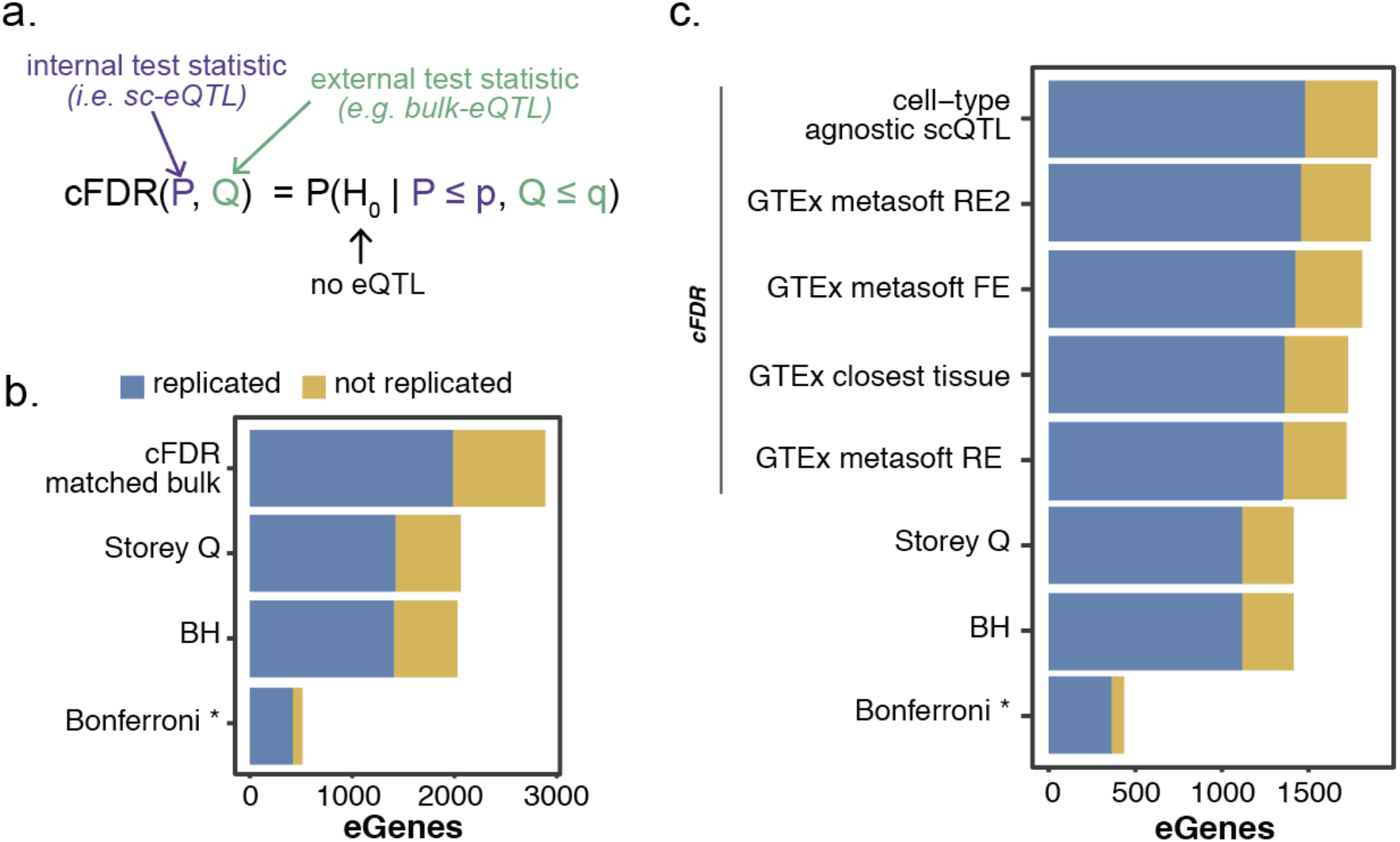
Conditional FDR increases eGene discovery in iPSC Smart-Seq2 data while replication fraction stays consistent. **(a)** Graphical summary of cFDR method. **(b)** The eGene discovery power using different FDR methods at an FDR of 5% for the respective method (p < 0.05 Bonferroni). Blue bars indicate eGenes that were replicated in all bulk (a-bulk) defined as FDR <10% and the same direction of effect. Yellow bars indicate eGenes that were not replicated in bulk. (number of total genes tested=20,334). **(c)** The eGene discovery power using different external test statistics for cFDR. Tests were performed only on the subset of genes present in all four external datasets (GTEx metasoft RE2, FE, RE, and closest tissue) (13,653 genes tested). * denotes multiple testing correction methods that control the (much stricter) family-wise error rate.

To assess the impact of using cFDR we used the dr-mean iPSC results and the eQTL statistics of the m-bulk samples to condition our FDR estimates on and used a-bulk samples to test the replication rates (**Fig. 5b, Table S8**). The cFDR eQTL results are compared against three “standard” multiple testing correction methods in eQTL mapping (Bonferroni [41], Benjamin-Hochberg [42] and Storey Q [43] -the latter being the standard method for our analyses [12,29,44–45]). We observed a 40% increase in eGene discovery by applying cFDR as compared to Storey Q, the second best performing method in terms of eGenes. When assessing the replication rates in a-bulk of the eQTL effects deemed significant by the different multiple testing procedures, we observe very similar replication rates for the methods seeking to control the false discovery rate. The Bonferroni method seeks to control the (much stricter) family-wise error rate, and as such has a higher replication rate but at the cost of making many fewer discoveries. Importantly, all eQTL identified by Storey Q are also significant in cFDR. In this specific case we could apply cFDR when only changing the expression data but keeping other settings matched, i.e. same donors and therefore the same genotypes. Because it is rare to have matched bulk and sc-RNAseq data for the same individuals, this is not a realistic setting for most applications of cFDR for sc-eQTL mapping. To generalise this approach we applied cFDR using other reference eQTL statistics to guide our sc-eQTL mapping analysis. Specifically, we selected two external reference sets from GTEx: 1) the GTEx tissue that most closely resembles iPSCs (EBV transformed lymphocytes [29]), and 2) the meta-tissue eQTL results by GTEx (v7). For the meta-tissue results we assessed all different meta-analysis p-values (i.e. the fixed effect, random effect and random effect 2 as calculated using metasoft [46]). Lastly, we used a cell-type agnostic eQTL mapping on the two other differentiation time points presented in the Cuomo et al study (i.e. mesendoderm and definitive endoderm). This final setting should be useful in most common sc-eQTL mapping settings where either a cell-type specific or cell-type agnostic analysis can be performed within the same study.

Similar to the results from the matched-sample cFDR, we observe that all of the cFDR methods (whichever test statistics are used for conditioning) outperform the Storey Q based multiple testing correction in terms of number of eGenes discovered (**Fig. 5c, Table S9**). Moreover, replication fractions are highly similar between the different FDR approaches (Storey Q and BH vs cFDR). The use of the p-values from the same study provided the largest increase in the number of eGenes, but the use of the metasoft p-values from the GTEx tissues is a close second (1,898 vs 1,859 eGenes). Of importance here is that we subsetted down to only consider genes tested in all datasets, to be able to compare between cFDR conditionings. When assessing the impact per reference p-value set, we see that leveraging the joint cell types from the Cuomo *et al*. study yields a comparable number of discoveries as using the m-bulk data (2,821 vs 2,887 eGenes), again with comparable replication fractions (**Table S10)**.

### Optimised sc-eQTL mapping

To showcase the total increase in power possible when combining the findings outlined above, we directly compared the eGene discovery results from a default workflow using d-mean aggregation (baseline) to dr-mean with an additional random effect to capture sampling variation and an improved multiple testing strategy applied. Again, we focused here on the iPSC data to be able to assess replication in bulk. Overall, we observe a 142.7% increase in eGenes detected when comparing the d-mean aggregation with Storey Q based multiple testing correction (1,402 eGenes) versus the optimised eQTL mapping approach, i.e. dr-mean with additional random effect and cFDR multiple testing (3,402 eGenes, **Fig. 6, Table S11**). This drastic increase is explained by the smaller individual optimisations discussed above. The switch to dr-mean instead of d-mean increases mapping power by 46.6%, the use of the additional random effect reflecting SV (1/#cells) increases power by an additional 31.9%, and lastly the guided multiple testing correction using p-values from the matched bulk analysis increases power by 25.5%. In terms of how well these sc-eGenes replicated the bulk-eGenes (using FDR<10% and same effect direction as replication criteria), we observe decreases in the fraction of sc-eGenes that replicated bulk-eGenes at each of the optimisation steps: ∼7% replication decrease for d-mean to dr-mean and an additional ∼4% replication decrease for the additional random effect on sample variation. However, the total number of replicated bulk-eQTL effects increased at every step, with 1,146 (+107.4%) additional eQTL replicated with the optimised mapping compared to baseline. When using a more stringent multiple testing cutoff (FDR<1%) in the optimal mapping procedure we observed a replication rate of 75.5%, which is similar to the replication rate for d-mean eQTL effects with Storey Q multiple testing (i.e. baseline), while still identifying 548 (51%) additional eGenes (**Table S12**).

**Fig. 6.**
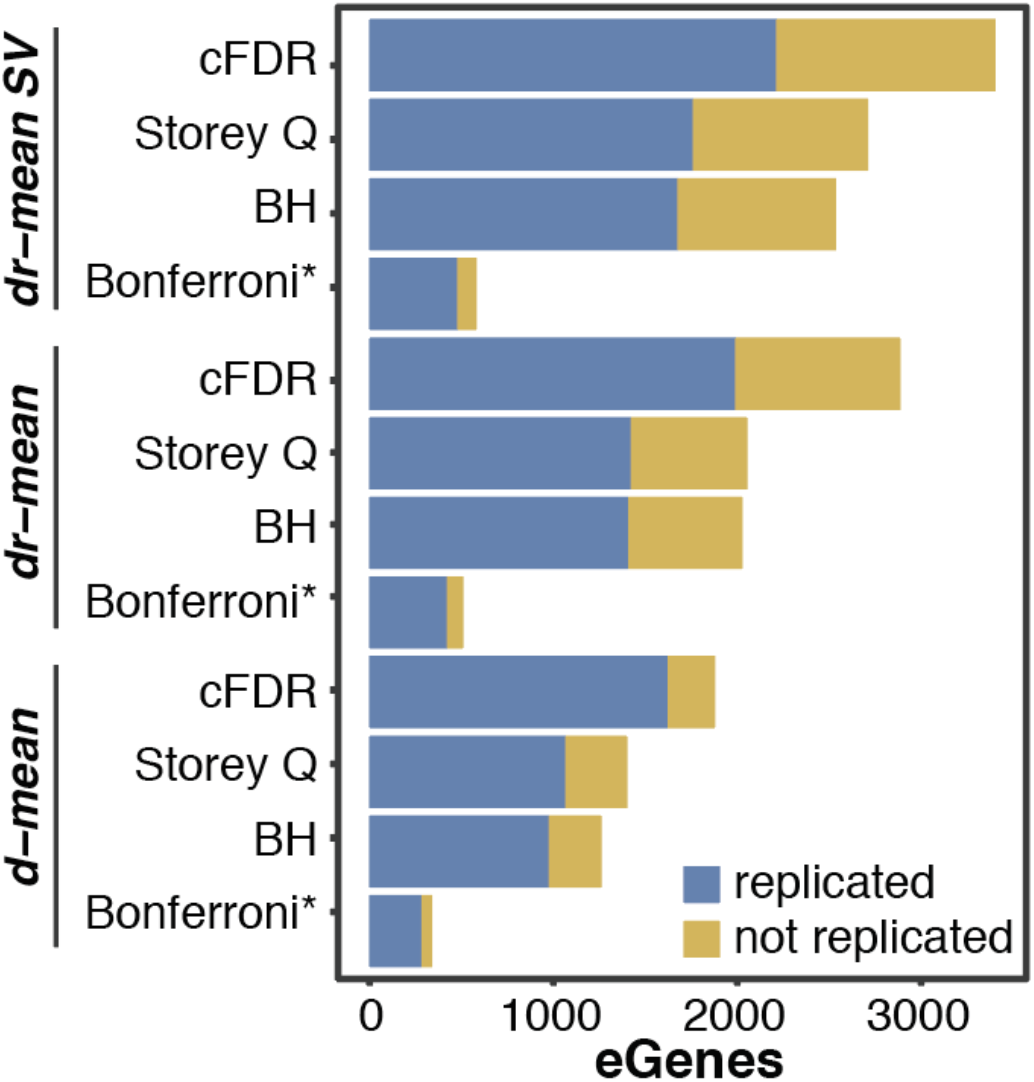
Optimising the eQTL mapping workflow increases eGene discovery. The eGene discovery power after optimising for aggregation level, including random effect reflecting sampling variation (SV, 1/#cells), and applying cFDR methods (conditioning on matched bulk p-values) at an FDR of 5% for the respective method (p < 0.05 Bonferroni). Blue bars indicate eGenes that were replicated in all bulk (a-bulk) defined as FDR <10% and the same sign. Yellow bars indicate eGenes that were not replicated in bulk (genes tested=20,334). * denotes multiple testing correction methods that control the (much stricter) family-wise error rate.

## Discussion

Since being highlighted as ‘Method of the Year’ in 2013 [47], sequencing of the genetic material of single cells has become common practice to investigate cell-to-cell heterogeneity in biological systems [48]. Now an established method, scRNA-seq is being applied on larger and larger scales, for example to chart all the cell types in the human body (Human Cell Atlas, HCA, [49]) and to study the differences between individuals from population-scale cohorts. This latter development has enabled us to study the effects of genetic variation on gene expression, at single-cell resolution. Recent studies have explored the optimisation of eQTL discovery power as a function of budgetary constraints, however, limited attention has been paid so far to optimising statistical workflows for eQTL mapping to improve power for the mapping of eQTL with the selected design.

Here, we set out to assess the effects of different eQTL mapping workflows for single-cell studies. We used bulk and single-cell expression profiles derived from the same iPSC lines to assess both power and replication of genetic effects on gene expression variation. We also tested both Smart-Seq2 and 10X data to determine how our recommendations would hold across technologies. To support our empirical data, we used splatPop to simulate data with known eQTL effects.

We find that optimising the aggregation and normalisation approaches can increase eQTL mapping power substantially: in the Smart-Seq2 study we identify close to two times more eGenes for the best performing method vs a non-optimised baseline method. In the 10X study, we find that the dr-mean approach also gives most power to identify eGenes, but we could not assess the replication in bulk. However, the 10X simulations show that dr-level aggregation results in fewer false discoveries, and that both mean and sum aggregation perform well. We speculate that whilst the sum would perhaps be the most obvious approach to reproduce bulk-like measurements, the mean might perform better because of the cell-level normalisation used. Indeed, the normalisation at the cell-level may better balance the differences in read counts across cells prior to the aggregation at individual-level. Additionally, it is of note that methods in which we treat replicates as separate samples (i.e. donor-run) work best even though only 20 (23%) of the Smart-Seq2 study samples and 25 (14%) for the 10X study samples were assayed in more than one run. We expect that these improvements are also down to improved normalisation as well as improved covariate correction. Finally, we tested median aggregation, but given the sparsity of the single-cell data this method drastically decreases the number of genes that can be assessed for eQTL mapping. Therefore, the use of median aggregation is not a sensible method in most single-cell applications.

Next we assessed the effects of both covariate and sampling variation correction approaches. First, the effect of the different methods to identify covariates, i.e. data driven factors that capture unwanted variation in eQTL mapping, is ambiguous. Adding covariate factors of any method increases eGene discovery power substantially, but the results for the different tools when using the optimal number of factors for that tool are broadly comparable both in terms of discovery and number of replicated effects. Correcting for 15 PCs yielded the most eGenes at a high replication rate. Importantly, the bulk eQTL results we use to assess replication were mapped after correcting the bulk RNA-seq data using 20 PCs, which might bias the replication for the sc-eQTL mapping to be more similar to the bulk PCA based results. However, regardless of the replication rate in bulk, PCA-correction yielded the highest number of eGenes and so remains a sensible “default” choice for defining covariates capturing unwanted variation. Second, including sampling variation as a random effect in the models results in an increase in eGene discovery power on both Smart-Seq2 and 10X datasets, especially when using the 1/#cells-based random effect.

Finally, we explored the use of bulk eQTL mapping statistics to guide our multiple testing correction using the conditional false discovery rate (cFDR) in the implementation by Liley et al. We observe that the cFDR method gives a marked increase in the identified eGenes (between 21% and 40% increase), both when matched bulk data is available and when meta-tissue eQTL summary statistics are used. However, using the larger iPSC dataset we found cFDR had limited impact in terms of replication rates of the eQTL. While cFDR allows us to guide the multiple testing, potential cell type-specific effects that are borderline significant might not be “boosted” in power by cFDR if those cell types are not present or lowly represented in the bulk references. We did, however, observe that all the effects found using standard FDR methods (Storey Q, Benjamini Hochberg) were also deemed significant using cFDR, so no potential effects were lost by using the guided FDR approach. We explored the use of cFDR for other single-cell studies, but more tests are needed for mixed cell types to find the optimal way to use cFDR in sc-eQTL studies.

This study demonstrates that optimising the sc-eQTL mapping workflow can increase eGene discovery power substantially. However, our conclusions come with some caveats; 1) simply discovering more eGenes does not necessarily mean that an approach is better, as false positives could arise due to data processing decisions; 2) bulk eQTL are a powerful, but not perfect, gold standard for assessing truth, as biases in bulk-eQTL mapping may be replicated in sc-eQTL mapping analyses; 3) while our simulation framework uses parameters estimated from empirical data to resemble real population-scale scRNA-seq data, all simulation frameworks have limitations. Also, the methods we explored here are based on current standards for bulk-eQTL mapping. Though we find that directly using single-cell expression information does not yield trustworthy sc-eQTL there is room for methodological improvements to better use the replicated structure of single-cell expression data and low counts in each cell.

In this paper, we discussed the impact of various normalisation, aggregation and covariate correction procedures as well as multiple testing correction in the context of single-cell eQTL mapping power. Together our optimised sc-eQTL mapping workflow increases eGene discovery by more than a factor two in our empirical data. These findings will be of great value to the community as more population-scale scRNA-sequencing data becomes available and as groups like the single-cell eQTLgen consortium are establishing sc-QTL studies on a massive scale.

## Methods

### iPSC RNA-seq data

To optimise processing and workflows for eQTL mapping in single cells we focus on iPSC cells, as iPSCs have a homogeneous and stable expression profile [29] and because of the availability of both single-cell [12] and bulk expression [2, 29] data for iPSCs for the same donors through the HipSci consortium. Genotyping of the iPSC lines is described in the original study [2] expression quantification per technology is described below.

### Single-cell RNA-seq data processing

The Cuomo et al [12] study describes in detail the differentiation of iPSCs towards definitive endoderm using a three day differentiation protocol. Briefly, the iPSC lines were multiplexed into 24 experimental pools (4-6 lines in each pool) and single cells were sequenced using the Smart-Seq2 protocol [33]). Cells were assigned to their donor of origin using Cardelino [7]. Here, we focus on the day0 cells (i.e. iPSC stage) that were successfully assigned to a donor and passed donor-and cell-level QC described below.

First, we quantified gene expression levels in line with the bulk study. We mapped RNA-seq reads to the genome (hg19) using STAR mapping and used ENSEMBL v75 for annotation. Expression levels for each gene were counted using featureCount (subread v1.6.0 [50]). To remove outlier runs (i.e. pools), we 1) calculated the average correlation of each cell with all other cells (across all runs), then 2) for each run calculated the median of the resulting average cell-cell correlations, and finally 3) discarded runs with a median cell-cell correlation < 0.7, leaving 30 runs (out of 35, **Fig. S8a**). Then, to remove outlier donors, we recalculated the cell-cell correlations between cells from the remaining runs and discarded donor-run combinations with a median cell-cell correlation < 0.5, leaving 155 donor-run combinations (corresponding to 7,611 cells; **Fig. S8b**). Additionally, to remove further possible confounding effects, a small group of donors with monogenic diabetes (n=10) and four donors that were outliers in the genotype space (based on projection with the 1000G project data) were excluded.

After QC and selection the number of cells per donor varied from 5 to 383 iPSCs per donor (mean=84.8) and the number of reads per donor varied from 1 to 104 million (mean=42 million). For reference, the average number of reads per donor in our bulk data is 44 million, but the range is much smaller (24-98 million reads per person, **Fig. S1**).

The two other developmental stages from Cuomo *et al*., i.e. mesendoderm and definitive endoderm are used as a reference set for the cFDR analysis. For these two developmental stages and cell-QC, both as described in the original publication. However, gene expression quantification was performed in line with the single-cell iPSC data described above.

### Bulk RNA-seq data processing

The HipSci project generated both chip-genotype and deep bulk RNA-sequencing profiles for 810 iPSC lines corresponding to 527 unique donors. Details on expression quantification and QC of these data can be found in [29]. Of the 111 pre-QC donors for which we had single-cell RNA-seq data, matching bulk iPSC data was available for 108 (97%). After QC and selection on both the bulk data and the single-cell expression data, we were left with data from 87 donors.

### 10X data from floor plate progenitor cells

Recent large-scale single-cell expression studies have mostly been conducted using the 10X Chromium platform [32], given its larger throughput in terms of number of cells and lower costs per cell. The quantification of expression from 10X is different from Smart-Seq2, because 10X sequencing 1) only aims to capture sequence from the 3’, 5’, or both 3’ and 5’ ends of transcripts and 2) uses unique molecular identifiers to identify and remove effects of PCR duplication, and 3) results in fewer reads per cell compared to Smart-Seq2.

Given these differences and the recent shift towards 10X data, we wanted to assess the impact of eQTL workflows on 10X data specifically. We selected one cell-type (midbrain floor plate progenitors; FPP) and time-point (day 11 of iPSC differentiation towards dopaminergic neurons) from Jerber et al [13], using expression quantification and cell QC from the original paper. Briefly, sequencing data generated from Chromium 10X Genomics libraries were processed using the CellRanger software and aligned to the GRCh37/hg19 reference genome. Counts were quantified using the CellRanger “count” command, with the Ensembl 84 reference transcriptome (32,738 genes) with default QC parameters. For each of 13 pooled experiments, donors (i.e. cell lines) were demultiplexed using demuxlet [9], using genotypes of common (MAF>1%) exonic variants available from the HipSci bank and a doublet prior of 0.05.

Sufficient single-cell data (>=5 cells) were available for 174 individual cell lines from 174 donors. The number of cells per donor ranged from 7 to 7,782 (mean=831.4). The total number of reads per donor ranged from 0.02 to 114 million reads (mean=9.7).

## Simulation Methods

Two types of population-scale scRNA-seq datasets (iPSC Smart-Seq2 and FPP 10X) were simulated using splatPop (http://www.bioconductor.org/packages/devel/bioc/vignettes/splatter/inst/doc/splatPop.html) from splatter [51]. To ensure realistic simulations, parameters were estimated from real data. Single-cell parameters defining the distribution of library sizes (gamma), common BCV (inverse chi-squared), and dropout rates (logistic) were estimated from the empirical single-cell count data from the donor with the most cells (iPSC Smart-Seq2 donor ID=joxm; FPP 10X donor ID=mita_1)s. Population parameters defining the distributions of gene mean expression (gamma) and variance in expression (gamma binned by mean expression) were estimated from the mean aggregated expression levels for all genes from either iPSC Smart-Seq2 or FPP 10X. Finally, eQTL parameters defining the distribution of the eQTL effect sizes (gamma) were estimated from the most significant eQTL hit for each gene from bulk iPSC data for eSNPs with MAF > 0.1. The eQTL effects were assigned randomly to 35% (n=439) of the genes being simulated with eSNPs having a MAF > 0.1 and being within 100kb of their eGenes. Data was simulated for genes from chromosome 2 (n=1,255) for 87 individuals in the British cohort for the Smart-Seq2 simulations and for 173 individuals from the European cohorts from 1000Genomes [52]. For the sample size analysis, n individuals were randomly selected from the European cohorts for each simulation (n=50, 87, 100, 173, 250, 400).

To mimic the experimental design of the empirical datasets, cells were simulated in batches with 5 (iPSC Smart-Seq2) or 24 (FPP 10X) individuals per batch, where 28% (iPSC Smart-Seq2) or 14% (FPP 10X) of individuals were replicated in two batches, reflecting the pooling sizes and replication rates in the empirical data. Batch effect sizes were sampled from log-Normal distributions with location=0.001 and scale=0.12 (iPSC and FPP) and with the splatPop parameter similarity.scale=5 (iPSC Smart-Seq2) or similarity.scale=22 (FPP 10X) set to reflect the batch effects and relationship between samples observed in the real data (**Fig. S2**).

The different eQTL mapping methods tested on simulated data were replicated 10 times, with each replicate being an independent splatPop simulation. Due to genetic linkage, the top eSNP mapped for each eGene was not expected to be the simulated eSNP. To account for this, all eQTL hits with an empirical p-value below the empirical p-value corresponding to the multiple testing threshold determined using Storey Q on the top hits for each gene were considered significant. Each simulated eGene was considered a true positive (TP) if the simulated eSNP was among the significant eSNPs or it was considered a false negative (FN). Note that the FN includes eGenes with no significant eSNPs (type 1) and eGenes with significant eSNPs, but for which the correct eSNP was not significant (type 2). Each simulated gene not assigned an eQTL effect was considered a true negative (TN) if it had no significant eSNPs and a false positive (FP) if it had a significant hit. Performance metrics include power (TP / (TP + FN)), empirical FDR (FP / (FP + TP)), and the Pearson’s correlation between simulated eQTL effect sizes and estimated effect sizes (beta.cor; including only genes simulated as eGenes). Statistical tests were performed in Rv4.0.3 and results are described in detail in **Table S3, S4**).

### Aggregation & normalisation methods

Single-cell normalisation and log2 transformation of normalised counts-per-million were performed on raw single-cell counts using scran/scater [25] with size factors calculated using the pooled approach described in [26] and a pseudocount of one applied to the log2 transformation to avoid attempting to take the log of zero (i.e. log2(cpm+1)). The normalised counts were then aggregated using either the mean or the median (**Fig. 1**). We also considered an alternative single-cell normalisation technique, bayNorm [27], but observed minimal differences in normalised counts (as assessed by Pearson’s correlation of at the gene-level over the individual cells) and in eQTL results after dr-mean aggregation (as assessed by overlapping eQTL effect size and p-values). Sum aggregation was performed directly on the raw counts (**Fig. 1**), and followed by pseudobulk-like TMM normalisation [24] and log2-transformation, as implemented in edgeR [28].

In all cases, aggregation was performed at two levels of batch (**Table 1**). First, we aggregated all cells from each donor (i.e. d-mean, d-median, d-sum). In this setting, one sample corresponds to one donor (n = 87 in the iPSC Smart-Seq2 dataset, n = 174 in the FPP 10X data; samples with > 5 cells only). Next, we aggregated separately across donors and sequencing runs (dr-mean, dr-median, dr-sum). In this second setting, one sample is a unique donor-sequencing run combination (when considering samples with > 5 cells, n = 155 in the iPSC Smart-Seq2 data, n = 702 in the FPP 10X data).

### Highly variable genes

Highly variable genes (HVGs) were defined as the genes in the top two quartiles based on their squared coefficient of variation (CV^2 = variance / mean^2) calculated across all cells of each different cell-type. In this manner, we identified 21,592 HVGs for the iPSC Smart-Seq2 dataset, and 16,369 for the FPP 10X dataset.

### eQTL mapping strategy

For *cis*-eQTL mapping, we followed Cuomo *et al*. [12], and adopted a strategy similar to approaches commonly applied in conventional bulk eQTL analyses. We considered common variants (MAF >10% and Hardy-Weinberg equilibrium P<0.001) within 100kb up-and down-stream of the gene body. Association tests were performed using linear mixed models (LMMs), fit using LIMIX [30]. The values of all genes were quantile normalised (“gaussianised” in LIMIX) and the significance was tested using a likelihood ratio test (LRT).

### Covariates for sc-eQTL mapping

To adjust for experimental batch effects across samples, we included covariates as fixed effects in our LMM models to correct for both known and hidden sources of unwanted variation. These covariates (e.g. batch effects) usually affect the expression of many genes, and therefore are detectable in the principal components of expression. Furthermore, global batch effects are orthogonal to the effects of a single *cis* regulatory variant on the expression of one gene, thus accounting for global batch effects in our LMMs will not remove the signal for eQTL. Unless specified otherwise, in our sc-eQTL mapping we included the first 20 principal components calculated on the relevant expression value aggregation/normalisation in the model as fixed effect covariates. However, in order to assess the impact of variations in the types and numbers of covariates included, we also considered alternative methods, namely PEER [36], MOFA (in two flavours: with and without sparsity; [38]) and LDVAE [37].

All covariates (except LDVAE, see below) were calculated using the relevant aggregated expression values at the level of donor or donor-sequencing run, for all genes tested (63,678 genes). PCA was computed using the R function prcomp. PEER (https://github.com/PMBio/peer/) was run using the R implementation with the number of factors set to 30 and default parameters. MOFA v2 (https://biofam.github.io/MOFA2/) was run using the R implementation, for n=25 factors. Both default parameters (i.e. MOFA; spikeslab_factors = F, spikeslab_weights = T, ard_factors = F, ard_weights = T) and parameters with the sparsity constraints removed (i.e. MOFA non-sparse; spikeslab_weights = F, ard_weights = F) were used.

Finally we included a recent approach using variational auto-encoders: linearly decoded variational autoencoder (LDVAE; [37]). In contrast to the other methods, LDVAE works directly on the single-cell count data. We ran LDVAE following the tutorial (https://www.scvi-tools.org/en/stable/user_guide/notebooks/linear_decoder.html), but changed it to estimate 25 factors over 1,000 epochs with 400 warmup epochs. After training we extracted the LDVAE factors and aggregated the factors over the cells by taking the mean per donor or donor-run.

### Mixed effect models

In order to account for possible population sub-structure in the sample considered as well as, importantly, for repeated observations for the same donor (i.e. across runs, in the “dr” aggregation methods) we include in the linear mixed model a random effect modelling such structure (using a kinship matrix, K):

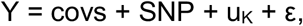

where Y is the expression of the gene considered (aggregated, normalised and transformed as described above), covs are the expression covariates (described above); SNP is the vector of allele values for the variant being tested; u_K_ is a random variable such that u_K_∼N(0, K), where K is a kinship matrix, specifically a SNP-based identity by descent (IBD) matrix estimated using PLINK [53], and ε is the error term.

Importantly, when considering “dr” aggregation methods (as well as the direct single cell eQTL mapping) this kinship matrix K is expanded to reflect the repeated samples (such that repeated measurements appear as “blocks” along the diagonal). Additionally, we tested the effect of the inclusion of a noise term in the LMM that accounts for the variable number of cells used to calculate aggregate expression measures, which has been shown to improve eQTL discovery power [13]. Here we also considered adding a term to account for each donor’s sequencing depth (i.e. number of reads per donor, **Table 4; Table S6)**. Using the LMM framework from LIMIX, we test three models for d-mean to account for different cofounders:

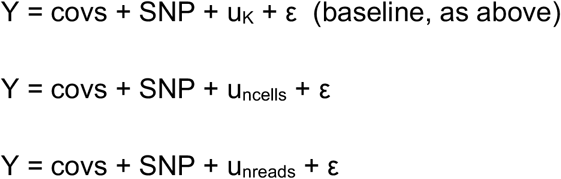

Where Y, covs, SNP, u_K_ and ε are defined as above; u_ncells_ is a random variable such that u_ncells_∼N(0, diag(1/ncells)), where diag(1/ncells) is a diagonal matrix with the inverse of the number of cells per donor as the diagonal terms and, analogously, u_nreads_∼N(0, diag(1/nreads)).

For dr-mean, since we have replicate measurements for each donor (across sequencing runs), we always need to account for population structure (and replicated structure), by including the (expanded) kinship matrix as a random effect. As such, we test the following models:

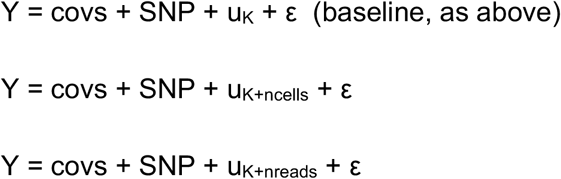

Since the LIMIX framework can only account for one random effect, in the latter two models, we introduced a weighting factor (*w*, between 0 and 1) to incorporate the relative weight of the two random effect term matrices, e.g. *w*\*1/ncells + (1-*w*)*K. We optimised *w* using a grid search per gene (Brent method, [54]), where *w* starts at 0.5 and is varied up or down in steps of 0.1 if the model fit increases with a higher or lower *w*.

## Multiple testing correction

Multiple testing correction for the eQTL results was performed in two steps, as is common-practice in eQTL studies [44, 45]. First, to adjust for multiple testing at the gene-level (i.e. across SNPs), we used an approximate permutation scheme, analogous to the approach proposed in [44]. Where, for each gene, we generate 1,000 permutations of the genotypes (100 permutations for simulated experiments) while keeping covariates, random effect terms, and expression values fixed. We then adjusted for gene-wise multiple testing using this empirical null distribution. Second, to control for multiple testing across genes, we applied the Storey Q value procedure based on the most significant eQTL per gene, unless otherwise specified. Genes with significant eQTL were reported at an FDR<5%. This second step is performed after gene selection (i.e. after considering the relevant gene selection in the comparison setting). We deem significant all associations that reach the gene-level corrected p-value (step 1) corresponding to the selected gene-level FDR (step 2), i.e. we call as significant all associations to a gene that are below the identified association p-value (not just the top associated SNP per gene).

We also tested alternatives to the Storey Q value for the second step. In particular, we test 1) the Bonferroni approach (p-value<0.05), which controls the much stricter family-wise error rate (FWER), 2) Benjamini-Hochberg (BH), another commonly used FDR approach;and 3) a recent implementation [39, 40] of the conditional FDR (cFDR), which leverages external data to guide the gene-level multiple testing correction. For the cFDR procedure we tested using raw association p-values from 1) the bulk iPSC associations, 2) GTEx v7 association p-values from the EBV transformed lymphocytes, 3) GTEx v7 association p-values from tissue meta-analyses results [55], and 4) association p-values from a joint eQTL mapping on the mesendoderm and endoderm data from the Cuomo et al study (Cuomo et al. 2020). Given that the external datasets have imperfect overlap in terms of both SNPs and genes, we limit our focus to overlapping genes and variants for each respective test, unless otherwise specified.

Conditional FDR is a method to condition or transform the p-value from one hypothesis test using an external set of test statistics. For a series of hypothesis tests (i = {1, .., n}), P = {P_i_} and Q = {Q_i_} are two sets of random variables, representing the observed p-values from the internal (i.e. sc-eQTL; p_i_) and external (e.g. bulk-eQTL; q_i_) hypothesis tests, respectively. Traditionally the cFDR is approximated with the empirical joint cumulative distribution function (see **Fig. 5a**) and is then used to draw a contour through the two-dimensional space of all p-value pairs (p, q)={(p_1_, q_1_), …(p_n_, q_n_)} that defines the region where H_0_ will be rejected. However, this approach does not explicitly control FDR. Here we apply an adjusted estimator of cFDR proposed and described in detail by Liley and Wallace. [40], which was shown to improve power and control the type-1 error rate. Briefly, for each p-value pair (p_i_, q_i_), the adjusted estimator adds randomly chosen points (p_i’_, q_i’_) to the original p-value pair space (p, q). Then the “transformed” p-value is calculated as the probability that the randomly chosen points had a more extreme cFDR than the point of interest (p_i_, q_i_). Ultimately, with this approach, we are able to define a better (i.e. a higher TP:FP) H_0_ rejection region than what would be defined by strict thresholds for P and Q, and thus better integrate the external information into the statistical test.

### Replication of bulk eQTL results

Performance of sc-eQTL mapping was measured using two metrics: number of significant sc-eQTL (FDR<5% unless stated otherwise, described above) and replication with bulk-eQTL. A sc-eQTL (i.e. eSNP-eGene pair with FDR<5% in a single-cell study) was considered to replicate a bulk-eQTL if it passed two criteria: 1) the gene-SNP pair was significant in the bulk test at FDR<10% and 2) the direction of the effect was consistent between the sc-eQTL and the bulk-eQTL.

Note that a small percentage (∼1%) of non-replicated sc-eQTL in bulk were due to the gene and/or the SNP of interest not being assessed in the bulk eQTL results.

## Acknowledgements

We would like to thank the sc-eQTLGen consortium for their input and useful discussions. We would like to thank Loukia Georgatou-Politou and PuXue Qiao for their help on applying cFDR in the context of single-cell eQTL mapping. We thank Oliver Stegle, Joseph Powell, Monique van der Wijst and Urmo Võsa for useful discussion, comments and edits to the manuscript.

## Author contributions

AC, GA, and MJB conducted eQTL mapping analyses with empirical bulk and single-cell RNA-seq data. CBA conducted the simulation studies. DJM and MJB supervised the work. AC, CBA, DJM, and MJB worked on the presentation and interpretation of results, and wrote the paper. This work was conducted as part of the sc-eQTLGen consortium. All authors read and approved the final manuscript.

## Data and code availability

The eQTL mapping pipeline is available via: https://github.com/single-cell-genetics/limix_qtl. splatPop is available in splatter v 1.14.1 on Bioconductor. The description of how to implement splatPop is available in the package vignette (http://www.bioconductor.org/packages/devel/bioc/vignettes/splatter/inst/doc/splatPop.html).

The HipSci genotype information is made available per sub study and are available via: PRJEB11750, EGAS00001000866, EGAS00001002015, EGAS00001002016, EGAS00001002013, EGAS00001002014, EGAS00001002011, EGAS00001002012, EGAS00001002010, EGAS00001002009, EGAS00001002006, EGAS00001001272, EGAS00001002008, EGAS00001001730, EGAS00001001273, EGAS00001002007, EGAS00001002005, EGAS00001000867

The single-cell iPSC, mesendoderm and endoderm RNA-seq reads are available via: ERP016000, EGAS00001002278.

The bulk iPSC expression data is available per sub-study via: EGAS00001000529, EGAS00001000593, EGAS00001001137, EGAS00001001318, EGAS00001001727, EGAS00001001986, EGAS00001001987, EGAS00001001988, EGAS00001001989, EGAS00001001990, EGAS00001001991, EGAS00001001992, EGAS00001001993, EGAS00001001994, EGAS00001001995, EGAS00001001996, EGAS00001001997, ERP007111

The FPP 10X data is available as part of the data published with (Jerber et al) at:https://zenodo.org/record/4333872#.YEJXnC2l2L4. Specifically, we considered the day 11 object only, and subsetted to the cell type of interest: FPP (floor plate progenitors).

Finally, bulk and single-cell iPSC count data from this manuscript specifically are available at: https://zenodo.org/record/4585385.

