## Supplementary Figures & Tables for "Optimising expression quantitative trait locus mapping workflows for single-cell studies"

### Table of contents

**Table of contents** 2

**Supplementary Figures** 3

**Supplementary Tables** 11

### Supplementary Figures

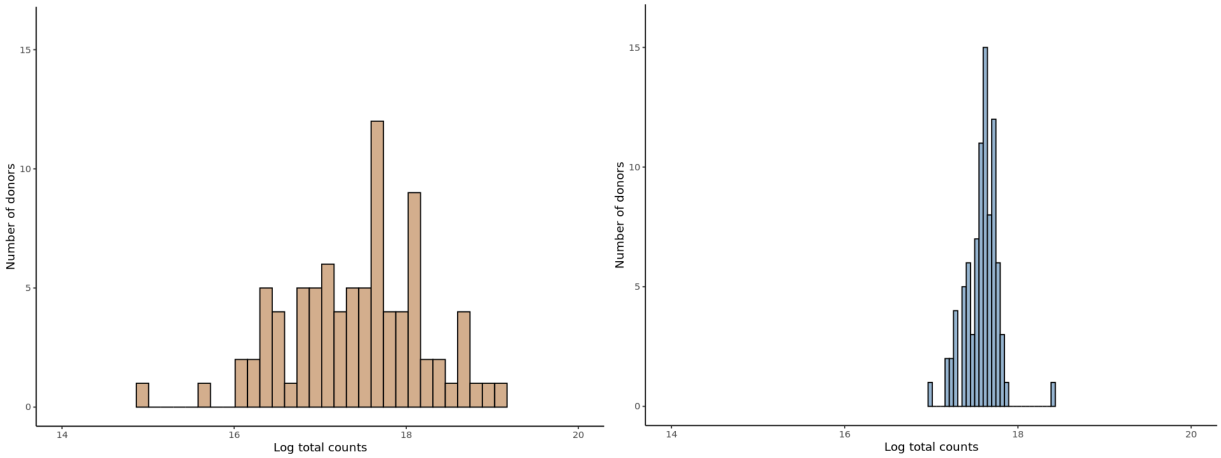


**Figure S1. Distribution of total reads between single cell and bulk RNA-seq data.** Distribution of total reads (genome-wide) per individual for matched iPSC data (from the same 87 individuals) using single cell SmartSeq2 (left) and bulk (right) RNA-seq data.

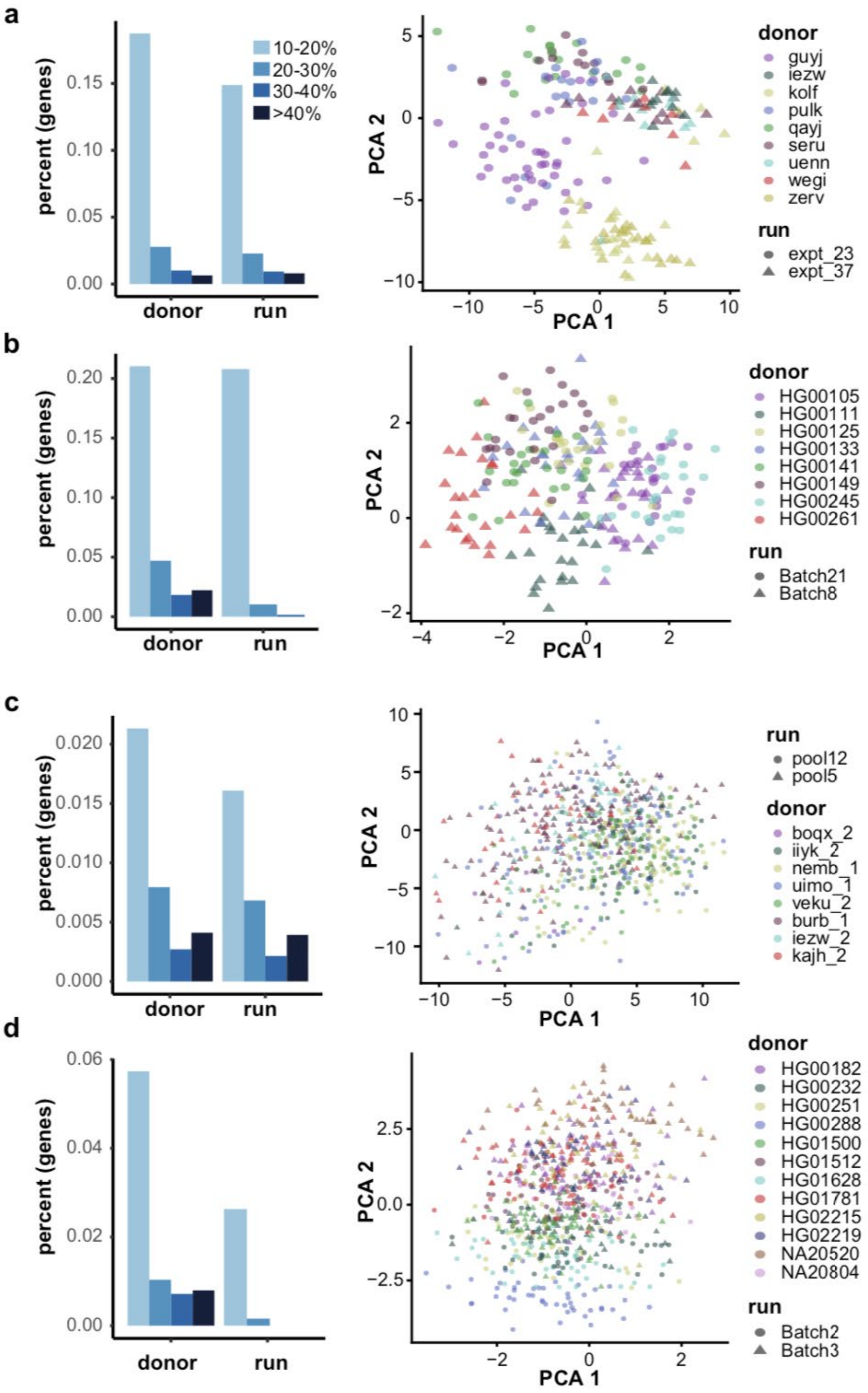


**Figure S2. Empirical and simulated population characteristics.** iPSC SmartSeq2 **(a)** empirical and **(b)** simulated data and NeuroSeq 10X **(c)** empirical and **(d)** simulated data. Barplots (left) show the percent of genes with differing degrees (color) of variance explained by the donor and run factors (x-axis). PCA plots (right) show a subset of cells for samples from two experimental runs, with cells shaped by run and coloured by donor.

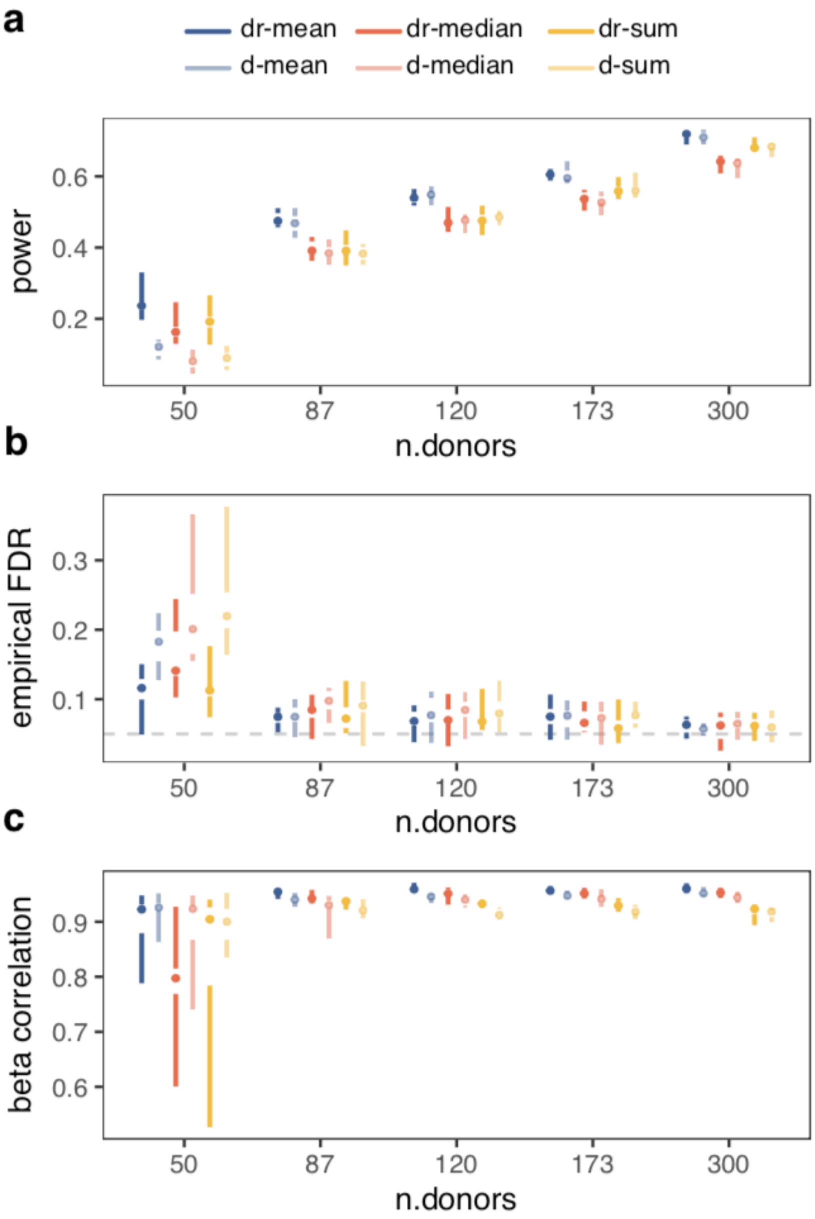


**Figure S3. Tufte's boxplots of eQTL mapping performance on simulated iPSC SmartSseq2 populations with the number of donors (x-axis) ranging from 50 to 300.** (**a**) Power to detect simulated eQTL (# true positives / # simulated eQTL). (**b**) Empirical FDR (false discovery rate at nominal FDR < 5%, dashed line). (**c**) Pearson's correlation between the ground truth and estimated effect sizes for genes simulated as eGenes. Colors as in Fig. 2. Shade designates aggregation level (donor-run: dark, left; donor: light, right). The point indicates the median, the gap indicates the interquartile range, and lines indicate the whiskers.

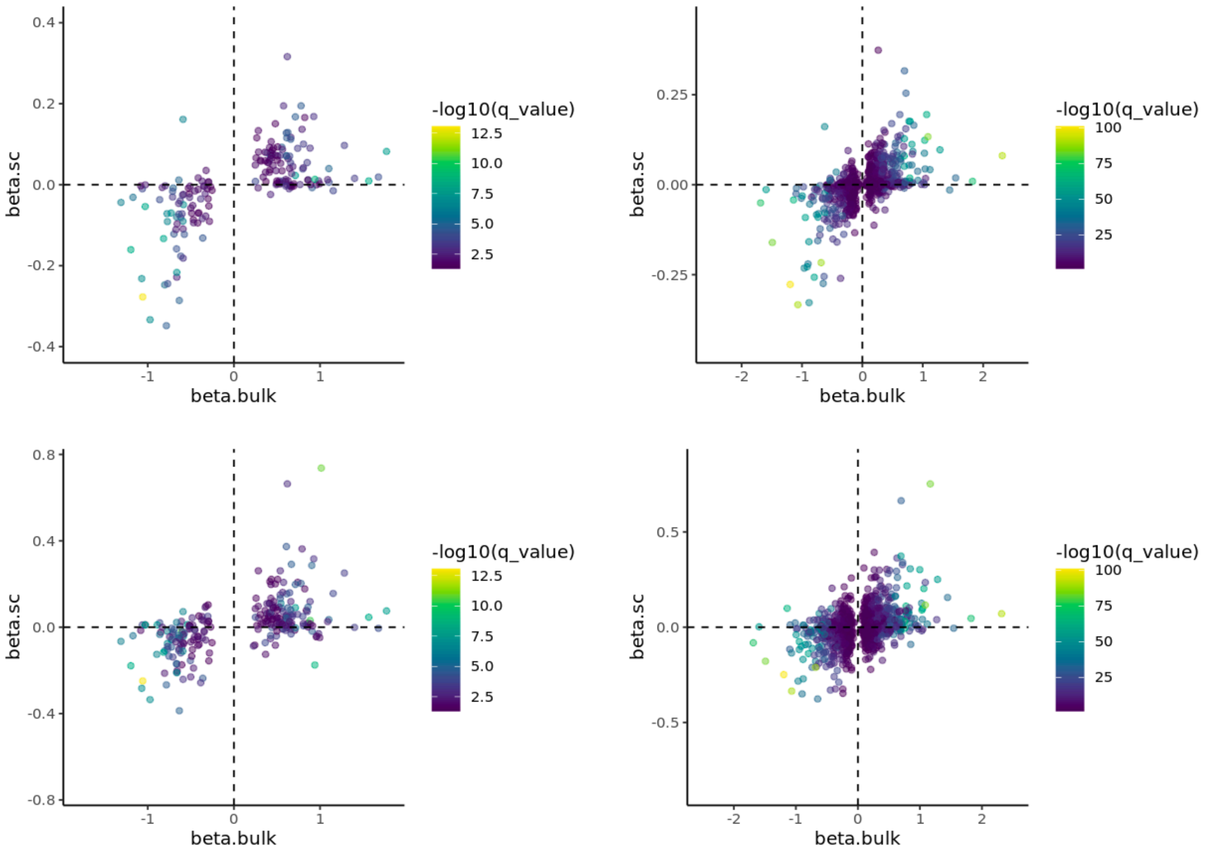


**Figure S4. eQTL mapping results of mapping eQTL directly on single-cell expression data.** Comparison of effect sizes between single-cell (y-axis) and bulk (x-axis) eQTL mapping results, when using individual cells as observations. Top: using all cells, bottom using only 5 cells per donor. Left: m-bulk, right a-bulk. Points are coloured by the q-value in the bulk results.

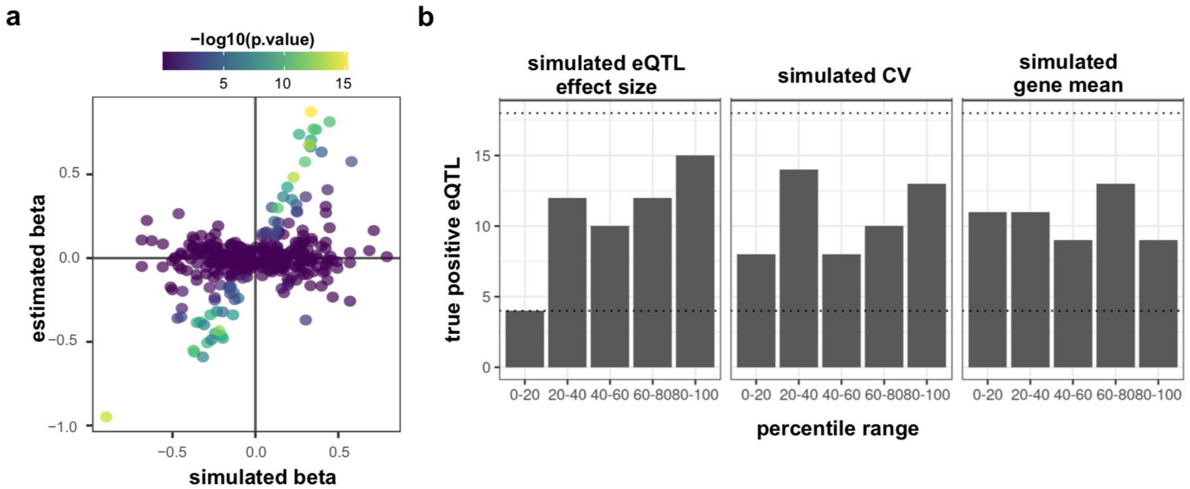


**Figure S5. eQTL mapping results of mapping eQTL directly on simulated single-cell expression data.** (a) Comparison of the estimated (y-axis) and simulated (x-axis) eQTL effect sizes for the 35% of genes simulated as eGenes from chromosome 2. Points are colored by the empirical p.value. (b) Number of true positive eQTL from each percentile range for simulated eQTL effect size, coefficient of variation, and gene mean. Dashed lines represent the 95th percentile range if we randomly sampled the same number of eQTL.

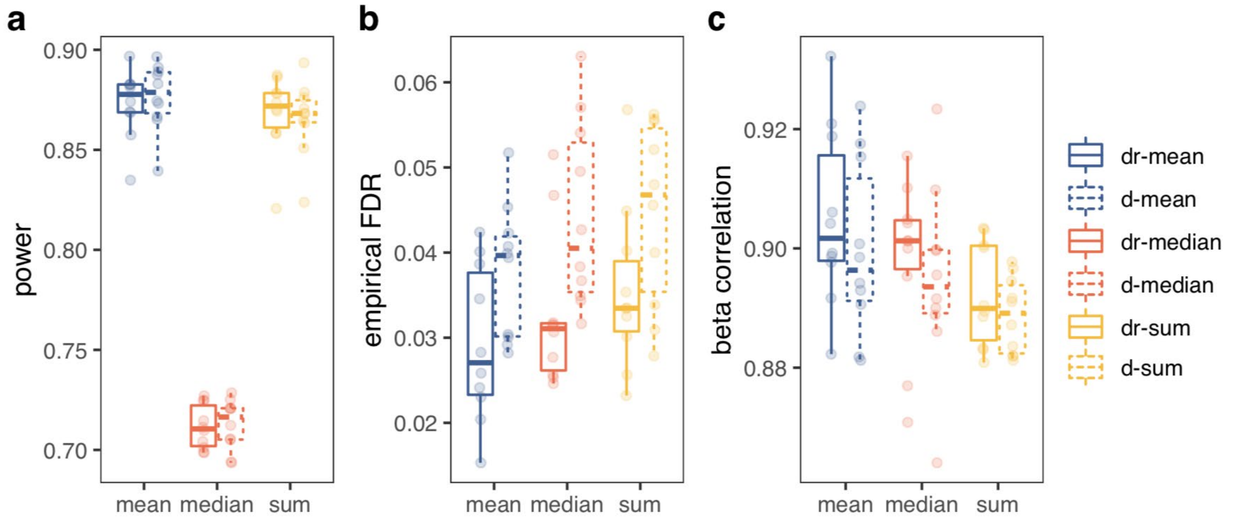


**Figure S6. Summary of eQTL mapping performance on simulated 10X neuron differentiation datasets. (a)** Power to detect simulated eQTL (# true positives / # simulated eQTL). **(b)** Empirical FDR (false discovery rate at FDR < 5%). **(c)** Pearson's correlation between the ground truth and estimated effect sizes for genes simulated as eGenes. Colors and line types are as in Fig. 2. Box plots summarise the distribution, while the points show performance for each replicate (n=10).

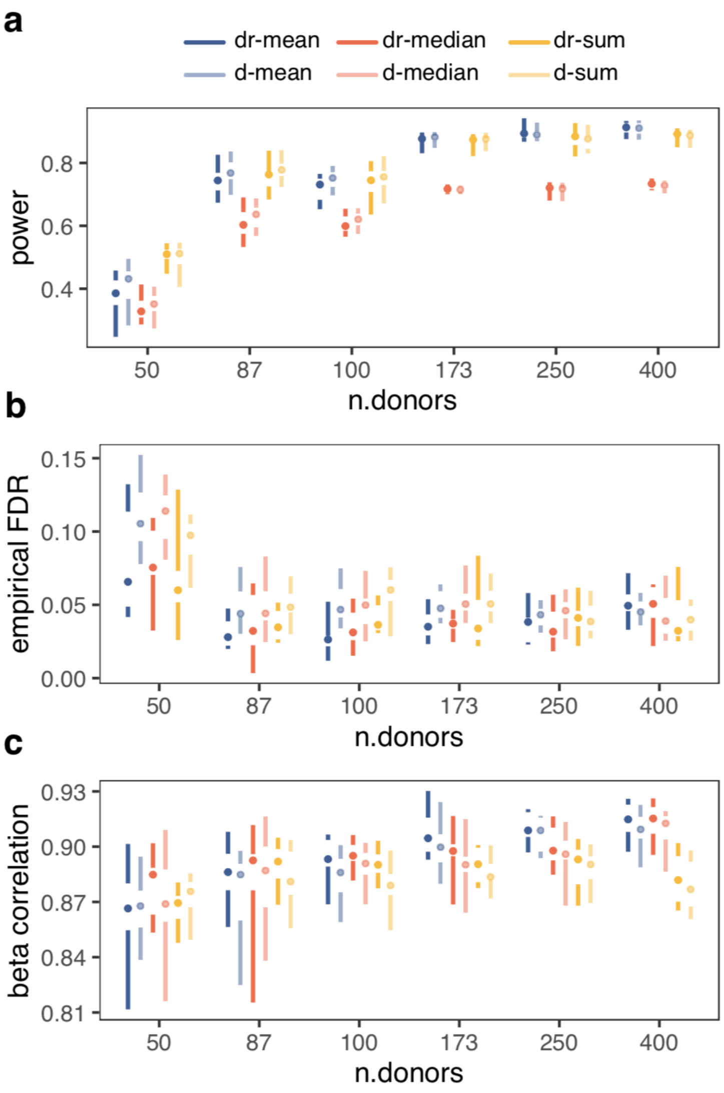


**Figure S7. Tufte's boxplots of eQTL mapping performance on simulated 10X neuron differentiation populations with the number of donors (x-axis) ranging from 50 to 400.** (**a**) Power to detect simulated eQTL (# true positives / # simulated eQTL). (**b**) Empirical FDR (false discovery rate at FDR < 5%). (**c**) Pearson's correlation between the ground truth and estimated effect sizes for genes simulated as eGenes. Colors as in Fig. 2. Shade designates aggregation level (donor-run: dark, left; donor: light, right). The point indicates the median, the gap indicates the interquartile range, and lines indicate the whiskers (n=10).

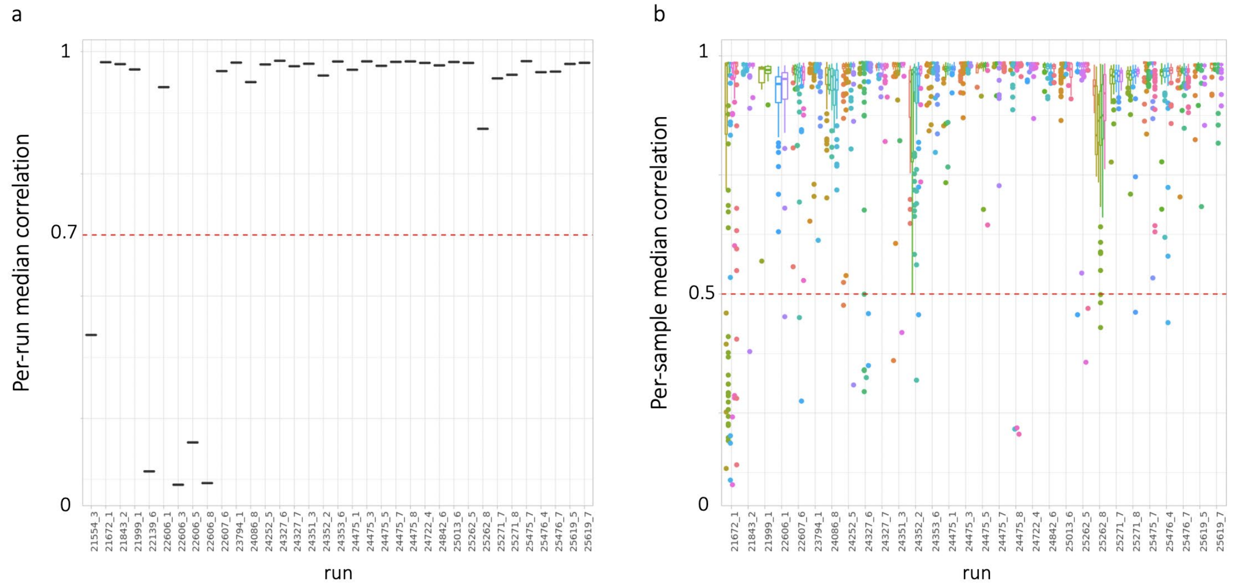


**Figure S8. Correlation-based cell QC.** (**a**) Median cell-cell correlation calculated per run. 1) For each cell, we calculated its average correlation with all other cells; 2) for each run, we calculated the median of these resulting average correlations; finally 3) runs with median cell-cell correlation < 0.7 (red dotted line) were discarded. (**b**) For the remaining runs, donor-run (i.e. sample) level median cell-cell correlations were calculated, and samples with median cell-correlation < 0.5 (red dotted line) were discarded.

### Supplementary Tables

**Table S1. Number of eGenes and replication of eQTL for the different aggregation & normalisation strategies in Smart-Seq2 iPSC cells.** Considering all highly variable genes (20,335 genes tested; **Methods**) tested (similar to **Table 1**). FDR was controlled at 5% for the discovery and we defined replication as FDR<10% and the same sign in both the matched bulk (N=87, m-bulk) and all bulk set (N=526, a-bulk).

| | **Discovery** | | **m-bulk replication** | | **a-bulk replication** | |
|---|---|---|---|---|---|---|
| | **eGenes** | **% tested** | **# replicated** | **% replicated** | **# replicated** | **% replicated** |
| **dr-mean** | 2,073 | 10.19% | 1,014 | 48.91% | 1,523 | 73.47% |
| **dr-sum** | 1,590 | 7.82% | 909 | 57.17% | 1,247 | 78.43% |
| **d-mean** | 1,415 | 6.96% | 854 | 60.35% | 1,123 | 79.36% |
| **d-sum** | 1,284 | 6.31% | 778 | 54.98% | 1,020 | 79.44% |
| **m-bulk** | 3,517 | 17.30% | - | - | 3,307 | 94.03% |

**Table S2 Simulation SmartSeq2 & 10X QTL mapping results.**
<External>

**Table S3 Detailed statistical test results from the SmartSeq2 simulation analysis.** P-values were adjusted for multiple testing when required using the bonferroni method. Significant tests (p or p.adj <=0.05) are in bold.

| Test | Effect | Grouped by | DFn | DFd | test.statistic | p | p.adj |
|---|---|---|---|---|---|---|---|
| **dependent variable: power** | | | | | | | |
| repeat measures 2-way ANOVA | level | | 1 | 9 | 9.45 | **0.013** | |
| | agg.method | | 2 | 18 | 262.41 | ***4.85e-14*** | |
| | level:agg.method | | 2 | 18 | 0.022 | *0.979* | |
| pairwise t-test | dr vs. d | | 29 | | 0.918 | *0.366* | |
| pairwise t-test | mean vs. median | | 19 | | 11.37 | 6.37e-10 | **1.91e-9** |
| | mean vs. sum | | 19 | | 13.77 | 2.43e-11 | ***7.29e-11*** |
| | median vs. sum | | 19 | | 0.57 | 0.574 | 1 |
| **dependent variable: empirical FDR** | | | | | | | |
| repeat measures 2-way ANOVA | level | | 1 | 9 | 15.0467 | ***0.004*** | |
| | agg.method | | 2 | 18 | 3.028 | 0.074 | |
| | level:agg.method | | 2 | 18 | 0.968 | 0.339 | |
| pairwise t-test | dr vs. d | | 29 | | -2.19 | ***0.036*** | |
| **dependent variable: beta correlation** | | | | | | | |
| repeat measures 2-way ANOVA | level | | 1 | 9 | 31.386 | ***3.3e-4*** | |
| | agg.method | | 2 | 18 | 16.346 | ***0.001*** | |
| | level:agg.method | | 2 | 18 | 0.154 | 0.858 | |
| pairwise t-test | dr vs. d | | 29 | | 3.824 | ***6.43e-4*** | |
| pairwise t-test | mean vs. median | | 19 | | 2.714 | 0.014 | **0.041** |
| | mean vs. sum | | 19 | | 6.612 | 2.5e-6 | ***7.5e-6*** |
| | median vs. sum | | 19 | | 1.207 | 0.242 | 0.726 |

**Table S4 Number of eGenes for the different aggregation and normalisation strategies in 10X midbrain floor plate progenitor cells.** In total 10,598 genes (HVGs tested in all methods excluding median) were considered in all of the strategies, and gene-level FDR was controlled at 5%.

| | **eGenes** | **% genes tested** |
|---|---|---|
| dr-mean | 3,167 | 29.88% |
| dr-sum | 2,516 | 23.74% |
| d-mean | 2,660 | 25.10% |
| d-sum | 1,939 | 18.30% |

**Table S5 Detailed statistical test results from the 10x simulation analysis.** P-values were adjusted for multiple testing when required using the bonferroni method. Significant tests (p or p.adj <=0.05) are in bold.

| Test | Effect | DFn | DFd | test.statistic | p | p.adj |
|---|---|---|---|---|---|---|
| **dependent variable: power** | | | | | | |
| repeat measures 2-way ANOVA | level | 1 | 9 | 0.498 | ***0.005*** | |
| | agg.method | 1.27 | 11.47 | 556.892 | ***1.87E-10*** | |
| | level:agg.method | 2 | 18 | 1.949 | 0.17 | |
| pairwise t-test | dr vs. d | 29 | | -0.0428 | 0.966 | |
| pairwise t-test | mean vs. median | 19 | | 44.7 | 1.04E-20 | ***3.12E-20*** |
| | mean vs. sum | 19 | | 1.49 | 0.15 | 0.46 |
| | median vs. sum | 19 | | -27.6 | 8.59E-17 | ***2.58E-16*** |
| **dependent variable: empirical FDR** | | | | | | |
| repeat measures 2-way ANOVA | level | 1 | 9 | 29.376 | ***4.22E-04*** | |
| | agg.method | 2 | 18 | 2.231 | *0.136* | |
| | level:agg.method | 2 | 18 | 0.187 | 0.982 | |
| pairwise t-test | dr vs. d | 29 | | -3.467 | ***0.002*** | |
| **dependent variable: beta correlation** | | | | | | |
| repeat measures 2-way ANOVA | level | 1 | 9 | 22.022 | ***0.001*** | |
| | agg.method | 2 | 18 | 3.927 | ***0.038*** | |
| | level:agg.method | 2 | 18 | 0.159 | 0.854 | |
| pairwise t-test | dr vs. d | 29 | | 1.06 | 0.297 | |
| pairwise t-test | mean vs. median | 19 | | 1.52 | 0.145 | 0.435 |
| | mean vs. sum | 19 | | 2.83 | 0.011 | ***0.032*** |
| | median vs. sum | 19 | | 1.61 | 0.124 | 0.372 |

**Table S6. Increased power in cis-eQTL mapping by correcting for covariates.** Reported are the percentages of number of eGenes (FDR<5%) out of 20,545 genes tested. Limited differences in terms of fraction of eGenes are observed. Highest number of eGenes is found when correcting for 15 PCs in this specific setting and test range.

| | 5 | 10 | 15 | 20 | 25 |
|---|---|---|---|---|---|
| **PCA** | 8.50% | 9.76% | 10.54% | 10.11% | 9.49% |
| **MOFA sparse** | 6.36% | 8.66% | 8.46% | 8.24% | 8.47% |
| **MOFA non sparse** | 7.96% | 7.67% | 8.06% | 8.11% | 5.07% |
| **PEER** | 8.48% | 9.54% | 10.36% | 10.14% | 10.45% |
| **linear scVI** | 6.28% | 7.07% | 7.28% | 7.72% | 7.96% |

**Table S7**. **Inclusion of random effect to increase discovery power of sc-eQTL mapping in 10X midbrain floor plate progenitor cells**. Shown are the number of eGenes that are discovered at an FDR of 5%. Tested are highly variable genes matched between the two considered aggregations (d-mean and dr-mean, n=10,598).

| | **Random effect matrix** | | | **Discovery** | |
|---|---|---|---|---|---|
| | **kinship** | **1/#cells** | **1/#reads** | **eGenes** | **% tested** |
| **donor mean** | ✓ | - | - | 2660 | 25.10% |
| **donor mean** | - | ✓ | - | 2731 | 25.77% |
| **donor mean** | - | - | ✓ | 2737 | 25.83% |
| **mean** | ✓ | - | - | 3167 | 29.88% |
| **mean** | ✓ | ✓ | - | 3140 | 29.63% |
| **mean** | ✓ | - | ✓ | 3317 | 31.30% |

**Table S8**. Conditional FDR increases eGene discovery in iPSC Smart-Seq2 data while replication fractions stay consistent. Shown are the number of eGenes that are discovered at an FDR of 5% of the resective method (p <0.05 for Bonferroni), and the replication in all bulk (a-bulk) defined as FDR <10% and same sign. Tested are all genes expressed when considering the mean aggregation/normalisation method and that are expressed in bulk (n=20,334).

| | Discovery (FDR 5%) | | all bulk all (FDR 10%) | |
|---|---|---|---|---|
| | **eGenes** | **% tested** | **# replicated** | **% replicated** |
| **cFDR** | 2887 | 14.20% | 1990 | 68.93% |
| **Storey Q** | 2058 | 10.12% | 1425 | 69.24% |
| **BH** | 2028 | 9.97% | 1410 | 69.53% |
| **Bonferroni** | 512 | 2.52% | 424 | 82.81% |

**Table S9.** Conditional FDR based on external datasets increases eQTL discovery power. Shown are the number of eGenes that are discovered at an (c)FDR of 5% of the respective method (p < 0.05 for Bonferroni), and the replication in all bulk (a-bulk) defined as FDR <10% and same sign. For comparison between the different reference sets we matched the genes between the different datasets considering only the genes that are expressed in all four datasets (n=13,653).

| | Discovery (FDR 5%) | | all bulk all (FDR 10%) | |
|---|---|---|---|---|
| | **eGenes** | **% tested** | **# replicated** | **% replicated** |
| **cFDR (cell type agnostic scQTL)** | 1898 | 13.90% | 1482 | 78.08% |
| **cFDR (GTEx metasoft RE2)** | 1859 | 13.62% | 1459 | 78.48% |
| **cFDR (GTEx metasoft FE)** | 1810 | 13.26% | 1426 | 78.78% |
| **cFDR (closest tissue GTEx)** | 1729 | 12.66% | 1364 | 78.89% |
| **cFDR (GTEx metasoft RE)** | 1720 | 12.60% | 1352 | 78.60% |
| **Storey Q** | 1413 | 10.35% | 1119 | 79.19% |
| **BH** | 1413 | 10.35% | 1119 | 79.19% |
| **Bonferroni** | 434 | 3.18% | 364 | 83.87% |

**Table S10.** Conditional FDR based on external datasets increases eQTL discovery power. Shown are the number of eGenes that are discovered at a cFDR of 5% when using the respective reference set (i.e. number of genes considered is different for every cFDR run), and the replication in all bulk (a-bulk) defined as FDR <10% and same sign.

| | | Discovery (FDR 5%) | | all bulk all (FDR 10%) | |
|---|---|---|---|---|---|
| | Genes tested | eGenes | % tested | # replicated | % replicated |
| **cFDR (cell type agnostic scQTL)** | 20540 | 2821 | 13.73% | 1852 | 65.65% |
| **cFDR (closest tissue GTEx)** | 16491 | 2183 | 13.24% | 1513 | 78.99% |
| **cFDR (GTEx metasoft RE2)** | 15550 | 2028 | 13.04% | 1602 | 78.99% |
| **cFDR (GTEx metasoft FE)** | 15550 | 1974 | 12.69% | 1567 | 79.38% |
| **cFDR (GTEx metasoft RE)** | 15550 | 1881 | 12.10% | 1489 | 79.16% |

**Table S11. Effects of different optimisation strategies and their combination on the power to map eQTL in single-cell data.** Shown are the number of eGenes that are discovered at FDR<5% of the respective method (FWER 0.05 for Bonferroni), and the replication in all bulk (a-bulk) defined as FDR<10% and same sign. Tested are genes expressed when considering the d-mean aggregation/normalisation method and that are expressed in bulk (n=20,334).

| **Settings** | | **Discovery (FDR 5%)** | | **all bulk all (FDR 10%)** | |
|---|---|---|---|---|---|
| **Mapping** | **Mult-test** | **eGenes** | **% tested** | **# replicated** | **% replicated** |
| **dr-mean SV** | cFDR | 3402 | 16.73% | 2213 | 65.05% |
| **dr-mean SV** | Storey Q | 2711 | 13.33% | 1760 | 64.92% |
| **dr-mean SV** | BH | 2536 | 12.47% | 1676 | 66.09% |
| **dr-mean SV** | Bonferroni | 582 | 2.86% | 479 | 82.30% |
| **dr-mean** | cFDR | 2887 | 14.20% | 1990 | 68.93% |
| **dr-mean** | Storey Q | 2055 | 10.11% | 1423 | 69.25% |
| **dr-mean** | BH | 2028 | 9.97% | 1410 | 69.53% |
| **dr-mean** | Bonferroni | 510 | 2.51% | 423 | 82.94% |
| **d-mean** | cFDR | 1879 | 9.24% | 1624 | 78.38% |
| **d-mean** | Storey Q | 1402 | 6.89% | 1067 | 76.11% |
| **d-mean** | BH | 1262 | 6.21% | 977 | 77.42% |
| **d-mean** | Bonferroni | 340 | 1.67% | 285 | 83.82% |

**Table S12. Effects of different optimisation strategies and their combination on the power to map eQTL in single-cell data.** Similar to Table 8, but considering FDR<1%**.** Shown are the number of eGenes that are discovered at an FDR of 1% of the respective method (P 0.05 for Bonferroni), and the replication in all bulk (a-bulk) defined as FDR <10% and same sign. Tested are genes expressed when considering the d-mean aggregation/normalisation method and that are expressed in bulk (n=20,334).

| Mapping | Mult-test | eGenes | % tested | # replicated | % replicated |
|---|---|---|---|---|---|
| **dr-mean SV** | **cFDR** | 2140 | 10.52% | 1615 | 75.47% |
| **dr-mean SV** | **Storey Q** | 1606 | 7.90% | 1195 | 74.41% |
| **dr-mean SV** | **BH** | 1550 | 7.62% | 1162 | 74.97% |
| **dr-mean SV** | **Bonferroni** | 473 | 2.33% | 396 | 83.72% |
| **dr-mean** | **cFDR** | 1798 | 8.84% | 1405 | 78.14% |
| **dr-mean** | **Storey Q** | 1283 | 6.31% | 985 | 76.77% |
| **dr-mean** | **BH** | 1274 | 6.27% | 981 | 77.00% |
| **dr-mean** | **Bonferroni** | 407 | 2.00% | 336 | 82.56% |
| **d-mean** | **cFDR** | 1284 | 6.31% | 1077 | 83.88% |
| **d-mean** | **Storey Q** | 893 | 4.39% | 716 | 80.18% |
| **d-mean** | **BH** | 829 | 4.08% | 666 | 80.34% |
| **d-mean** | **Bonferroni** | 251 | 1.23% | 205 | 81.67% |
